# Epigenomic Landscape of Oak (*Quercus robur*) across Seasons and Generations

**DOI:** 10.1101/2025.11.14.688509

**Authors:** Rosa Sanchez-Lucas, Joe He, Marcella Chirico, Jack L. Bosanquet, Annabella A. B. Lehner, Kehinde S. Oyesola, Kris Hart, Rob A. MacKenzie, Estrella Luna, Marco Catoni

**Affiliations:** School of Biosciences, University of Birmingham, Birmingham B15 2TT, UK; Birmingham Institute of Forest Research, University of Birmingham, Birmingham B15 2TT, UK; School of Geography, Earth and Environmental Sciences, University of Birmingham, Birmingham B15 2TT, UK

**Keywords:** DNA methylation, epigenetics, oak, forest trees, transposable elements, transgenerational memory, *Quercus robur*, Transgenerational effect, Long-lived plants, Seasonal regulation

## Abstract

- Seasonal fluctuations strongly shape the physiology of long-lived trees by coordinating growth, dormancy, and stress responses. Increasing evidence points to epigenetic mechanisms, particularly DNA methylation, as regulators of these processes, yet their role in long-lived trees across seasons and generations remains poorly understood.
- We generated single-base resolution maps of cytosine methylation exploring the epigenetic landscape of 180 year-old mature oak (*Quercus robur*) trees (genetically homogeneous) along spring, summer and autumn, and in their progeny.
- Genome-wide DNA-methylation revealed a progressive increase in the CHH context (H = A, T or C) from Spring to Summer and Autumn, suggesting epigenetic reprogramming is happening over season. Differentially Methylated Regions (DMRs) were concentrated in promoter regions and terminal inverted repeat (TIR). Differentially methylated transposable elements (TEs) and genes were involved in leaf development and hormonal signalling. By contrast, generational differences (parents versus offspring) were most prominent in CG and CHG contexts and were concentrated in genic regions.
- Oaks exhibit distinct seasonal and generational DNA methylation signatures, highlighting the plasticity and developmental specificity of epigenetic regulation. These findings provide a genomic foundation for understanding how epigenetic memory contributes to phenology, developmental programming and long-term adaptation in long-lived plants.

## Introduction

The Quercus genus, commonly known as oaks, comprises over 500 species distributed throughout Europe, the Americas, Asia, and North Africa. In northern and central Europe, *Quercus robur* (English oak) and *Q. petraea* (sessile oak) dominate native woodlands, where they are a sink for carbon sequestration and climate resilience (Bratu *et al*., 2025) and play critical ecological roles (Bölöni *et al*., 2021) while holding significant cultural and historical value (Haneca, 2009).

Trees, and oak in particular, undergo cyclic seasonal changes linked to the developmental transition from a vegetative phase to reproduction. These switches are genetically determined by the integration of endogenous factors (e.g., age and developmental state) and environmental inputs, such as day length, light quality, and temperature (Amasino, 2010; Andrés & Coupland, 2012; Song & Chen, 2015). In *Quercus robur*, spring triggers dormancy release, bud burst, and active photosynthesis, driven by increasing temperatures and photoperiods (Benito Garzón *et al*., 2024). Spring also marks the onset of flowering in mature trees, with male catkins typically emerging before or alongside leaf development, followed shortly by female flowers (Jato *et al*., 2015). Pollination occurs primarily via wind (Ashley, 2021), and fertilisation leads to the acorn development through the summer months (Miguel *et al*., 2015), during which they act as strong sinks for photoassimilates. Photosynthetic capacity increases over summer, reaching a peak by the end of the season, just before senescence (Burnett *et al*., 2021). By autumn, acorns reach maturity and begin to fall, coinciding with the onset of leaf senescence and nutrient reallocation before winter dormancy (Burnett *et al*., 2021). These seasonal transitions are tightly coordinated by hormonal signalling and metabolic reprogramming and are increasingly being linked to underlying epigenetic reconfigurations (Rudy *et al*., 2024; Silva *et al*., 2024), although the number of epigenetic analyses performed on forest trees are still very limited (Peck & Sork, 2024).

At the mechanistic level, epigenetic variation is primarily mediated by DNA methylation and histone modifications. Among these, DNA methylation is more readily and precisely measurable using genomic approaches, making it the most studied epigenetic mark in non-model species. In plants, DNA methylation occurs at cytosine bases within CG, CHG, and CHH contexts (with H indicating A, T or C) (Zhang *et al*., 2006; Zilberman *et al*., 2007). The symmetrical nature of CG and CHG sequences, maintained by the plant METHYLTRANSFERASE1 (MET1) and CHROMOMETHYLASE3 (CMT3) enzymes, allows for stable inheritance during cell division, as the methylated cytosine on the parental strand directs methylation on the newly synthesized strand (Bartee *et al*., 2001). In contrast, CHH methylation lacks symmetry, requiring 24 nucleotides (nt) sRNAs to guide the DOMAINS REARRANGED METHYLTRANSFERASE 2 (DRM2) in re-establishing methylation independently by replication with a process called RNA-directed DNA methylation (RdDM), mediated by the plant specific RNA polymerases PolIV and PolV (Cao & Jacobsen, 2002).

In oak, epigenetic research has shed light on a range of mechanisms of control in biological processes. For example, variation in the DNA methylation of valley oak (*Quercus lobata*) seedlings has been associated with plant height, leaf lobedness, powdery mildew infection, and trichome density (Browne *et al*., 2019). Analysis performed with the reduced-representation bisulphite sequencing approach identified specific single-methylation variants (SMVs) associated with climate variables, particularly temperature (Gugger *et al*., 2013). These SMVs were often located near genes involved in environmental response, indicating a potential role for DNA methylation in local adaptation (Gugger *et al*., 2013). In holm oak (*Quercus ilex*), a study encompassing adult, embryo, and seedling stages revealed that global DNA methylation levels were highest in adult organs, with embryos exhibiting more demethylation events and seedlings showing more de novo methylation events. These findings suggest that DNA methylation patterns are dynamic and stage-specific, influencing developmental processes (Labella-Ortega *et al*., 2024).

Over the past decade, high-quality genome assemblies have been generated for multiple oak species (Sork *et al*., 2016; López-Hidalgo *et al*., 2018; Ramos *et al*., 2018; Kapoor *et al*., 2023; Rey *et al*., 2023), including *Q. robur (*Plomion *et al., 2018)*. All sequenced oak genomes share a conserved chromosome number (n=12) and contain 20,000 to 40,000 protein-coding genes, offering a foundation for genomic research. These resources have facilitated comparative genomics and functional studies, shedding light on traits like drought tolerance (Madritsch *et al*., 2019), disease resistance (Conrad *et al*., 2019) and the production of secondary metabolites in cork (Jato *et al*., 2015; Han *et al*., 2022). However, how epigenetic landscapes in oaks are shaped by seasonal cycles and generational transitions remains largely unexplored. In this study we provide the most complete evaluation of the seasonal and generational epigenomic landscape of *Q. robur.* This work provides new insight into how the epigenome of a long-lived plant species is modulated by seasonal and developmental factors, laying a foundation for future research into the role of epigenetic mechanisms in long-term environmental responsiveness.

## Materials and Methods

### Plant Material and Experimental Design

Mature English oak (*Quercus robur*) trees (Table S1) were selected from the ambient-air arrays of the Birmingham Institute of Forest Research (BIFoR) Free Air CO₂ Enrichment (FACE) facility in Staffordshire, UK (Hart *et al*., 2020). Six trees (IDs: 8749, 8673, 8343, 9301, 6387, and 6382) were chosen for seasonal and generational epigenomic profiling. Leaf samples were collected from the mid-canopy of these trees during three distinct phenological stages: Spring (27th April 2021), Summer (29th June 2021), and Autumn (28th September 2021). Images of representative leaves at the different seasonal collections can be found in supplementary Figure 1 (Figure S1). Professional arborists conducted the collections, retrieving three to five leaves per tree per time point. Leaves were collected from the six trees in Spring, from two trees in Summer and from three trees in Autumn (Table S1). Upon harvest, samples were flash-frozen in a dry shipper (Statebourne Biotrek 10) and stored at -80°C until DNA extraction.

During the Autumn collection, acorns from the same trees were enclosed in mesh fruit bags (TDCuizent, 23EK19T4914U53Y8P69UW0040C2J6E) to prevent herbivory and ensure parental identity. Bags were retrieved on the 5th of November 2021 once acorns had reached maturity. Collected acorns were washed and pre-germinated as previously described (Sanchez-Lucas *et al*., 2023), then transferred to Maxi Rootrainers (Haxnicks, RT230101) filled with 400 mL compost (peat-free Levington Advanced F2). Seedlings were grown under controlled glasshouse conditions (16 h light/8 h dark; ∼200 µmol m⁻² s⁻¹; 20°C/18°C; 42% relative humidity) and watered to field capacity. Leaves from seedlings with at least four fully expanded leaves were collected and flash-frozen in liquid nitrogen. All samples were stored at -80°C until further processing. Samples coming from the same tree ID, including the offspring, were considered part of a genetically related group and were defined as a “Family” (Table S1).

### Diversity Arrays Technology sequencing

Leaf samples had previously been collected from the same six mature oak trees in July 2018, using a similar canopy sampling strategy but collecting material from the bottom, middle and top of the canopies. The three samples serve as technical replicates per tree. These samples were stored at -20°C prior to DNA extraction for genotyping-by-sequencing (GBS) analysis using the DArTseq method from Diversity Arrays Technology (Canberra, ACT, Australia) (Egea *et al*., 2017). As additional control, we included leaves samples from pure *Quercus robur* and pure *Q. petraea*, supplied by Forestar company using the UK provenance 403 for both species. Plants were grown in the same conditions as the offspring seedlings (described above). DNA was extracted from 100 mg of ground frozen leaves using the DNeasy Plant Mini Kit (Qiagen) following the manufacturer’s instructions with minor modifications and resuspended in 100μl elution buffer. Extracted DNA was digested using the restriction enzymes combination *PstI-MseI*, followed by adapter ligation to enrich polymorphic regions of the genome. Polymorphic regions were cloned in *E.coli* and libraries generated. Amplicons were size selected and sequenced. Sequences were aligned using the *Quercus robur* genome (Plomion *et al*., 2018) and SNPs and SilicoDArT markers were searched and filtered using DArT pipelines. Data resulted in presence/absence matrices (Kilian *et al*., 2012). Hierarchical clustering and dendrogram plot were performed using the Pearson correlation as the similarity measure and distances defined as 1-r, where r is the Pearson correlation coefficient; the clustering method employed was the average linkage method (UPGMA).

### Whole Genome Bisulfite Sequencing

Genomic DNA was extracted from approximately 100 mg of tissue from each sample, using the Qiagen DNeasy Plant Mini Kit (Qiagen) following the manufacturer’s instructions with minor modifications and resuspended in 100μl elution buffer. A total of 400 ng of Genomic DNA was used for Whole Genome Bisulfite Sequencing (WGBS), was performed by Novogene (Beijing, China) using the Illumina NovaSeq 6000 platform.

Raw WGBS sequence data for the 14 sequenced samples were obtained as FASTQ files and quality control and trimming were performed using fastp v0.23.2 (Chen *et al*., 2018) with default parameters. The reads were aligned with the reference oak genome (Plomion *et al*., 2018) (accessed 06/11/23) using Bismark v0.22.3 (Krueger & Andrews, 2011) with the “*--bowtie2*” v 2.3.5.1 (Langmead & Salzberg, 2012) alignment option. The output was deduplicated using “*deduplicate_bismark*” and cytosine methylation counts were extracted using “*bismark_methylation_extractor*” and classified in the three main context CG, CHG and CHH (H = A, T or C), generating a CX_report file per sample. PCAs were computed using the “*prcomp*” function from the *stats* R package for each cytosine context, starting from the called “CX_report file”.

### Differentially Methylation Regions (DMRs) Calling

To call Differentially Methylation Regions (DMRs), we employed the *DMRcaller* R package *(Catoni et al., 2018b; Catoni & Zabet, 2021)* using the function “*computeDMRsReplicates*” in each Group comparison separately for the three cytosine contexts, using the parameters: bin size of 400 nt (*binSize*), a cutoff for methylation difference of 0.2 (*methylationDiff*), a minimum count of 2 cytosines (*minCytosinesCount*) and a minimum of 4 reads per cytosine (*minReadsPerCytosine*). Overlapping DMRs were found using the “*overlapsAny*” function from the *GenomicRanges* package (Lawrence *et al*., 2013) in R and plotted using the *venn.diagram* package in R.

### Genetic Feature Overlaps

*ab initio* and manually curated gene features associated with the oak reference genome (Plomion *et al*., 2018) were used for genetic feature annotations, restricted to the 12 main chromosomes of oak. Genes were defined using the “mRNA” feature from the reference annotation and promoters were defined as being 1000 bp upstream of “mRNA” features, computed using the “*flank*” function from the *GenomicRanges* package. TEs were filtered to be >500 bp, of a known class and of the higher confidence “match” type for further analysis. Intergenic Regions were defined as regions that were not a gene, promoter or TE. A custom function randomly placed 10,000 regions (400 nucleotides each) in the reference genome to create “Random” DMRs, which were then compared to the actual observed distributions. Overlaps between DMRs and gene features were identified using the “*overlapsAny*” function from the *GenomicRanges* package (Lawrence *et al*., 2013). Proportions of overlaps with the different genetic elements were computed and displayed as stacked bar plots. Statistical significance between overlapping regions was done using the “*fisher.test”* function.

### Feature Methylation

For line graphs of methylation proportions, the appropriate features were first converted into bedgraph format using the “*export*” function from *rtracklayer* R package. *Deeptools* (v 3.5.2) package (Ramírez *et al*., 2014) was used to compute methylation matrices (standard parameter, except with the “computeMatrix” option, the –scale-regions flag and --binSize 200) for methylations for each bedgraph and the “*plotProfile*” function to plot results.

For the heatmaps, the methylation proportions of disaggregated samples were computed using the “*analyseReadsInsideRegionsForCondition*” function from *DMRCaller* in regions formed from the union of all 6 pairwise DMR calls of the Spring, Summer, Autumn and Offspring groups. This was repeated using the loci of filtered TEs and the DMRs called between Parents and Offspring, respectively. The resulting matrices were clustered and visualised using the *pheatmaps* R package, using default clustering parameters.

### Gene Ontology Enrichment

Ontology was performed using a previously described approach (Sanchez-Lucas *et al*., 2025) with annotation performed on the oak reference genome (Plomion *et al*., 2018) and terms identified using Trinotate v4.0.2. In total, 21710 transcripts were mapped to GO terms for further analysis. GO enrichment analysis was performed in R using *topGO* v2.54.0 on the gene features overlapping with DMRs (identified as described above). The Weight01 algorithm and Fisher’s exact test were used to determine significantly enriched GO terms. Statistically significant GO terms (p < 0.02) with at least 10 gene matches were identified in the CG, CHG and CHH contexts for the biological processes (BP) class.

### SNP Elucidation from Whole Genome Bisulfite Sequence

SNPs were identified using CGmapTools (v0.1.2) (Guo *et al*., 2017) with standard parameters via the Bayesian strategy using dynamic p-value. Trimming and quality control were carried out using Fastp (v0.23.4) (Chen *et al*., 2018) and FastQC (v0.12.1). Indexing, alignment, and mapping were performed using Picard (v2.27.4), Bismark (v0.24.2) (Krueger & Andrews, 2011), and Bowtie2 (v2.5.1) (Langmead & Salzberg, 2012), with further processing on the resulting BAM files (adding ReadGroups and sorting by co-ordinates) using Picard “*AddOrReplaceReadGroups*” and “*SortSam*”. This SNP calling pipeline was performed on all samples processed in parallel via Nextflow (v25.04.6). The resulting VCF output files were merged, G/C and A/T filtered by read depth and converted to BED format using PLINK (v1.90b6.24) (Purcell *et al*., 2007). Further filtering based on minor allele frequency (MAF) and principal component analysis were performed using the R (v4.3.2) package *SNPRelate* (v1.6.4) (Zheng *et al*., 2012) and visualised by using *ggplot2* (v3.5.2) R package.

### Sequence Frequency

We used the mappability score to measure the level of frequency of any DNA sequence in the oak genome, as previously described (Catoni *et al*., 2017). The mappability of all 20mers (mismatch = 1) were computed across the whole oak genome using GenMap (Pockrandt *et al*., 2020) and their frequency (inverse of the mappability) was computed. To investigate the frequency of the most differentially methylated TEs, the difference in methylation of Spring and other Seasons for each TE was computed and the 10% with the greatest and smallest difference was identified. A violin plot of their frequencies and medians was presented.

## Results

### Seasonal and Generational DNA Methylation Dynamics of Oak Trees

Prior to investigating epigenetic variation, we assessed the genetic background of the sampled trees at our study site. Hierarchical clustering based on SNP’s structural variants identified by DArTseq confirmed that all individuals belong to *Q. robur* and exhibit a high degree of genetic uniformity (Figure S2a), showing that our sampled oak individuals derived from a homogenous genetic pool. Then, epigenetic variation across seasons and generations was evaluated on green leaf material collected from oak trees in Spring, Summer and Autumn (Parents) of 2021, and their progeny (Offspring) (Figure 1a). The obtained genome-wide cytosine methylation profile at single base resolution for each sample were sorted into CG, CHG and CHH contexts. PCA analyses revealed that Summer and Autumn samples are grouped primarily by individual and family, likely reflecting genetic relationships among individuals (Figure 1b). However, a strong seasonal effect was evident in Spring samples, consistently separated from Summer and Autumn samples in all three cytosine contexts (Figure 1b), indicating distinct epigenomic states early in the growing season. This seasonal epigenetic divergence was especially pronounced in the CHH context, where Spring samples formed a clearly defined cluster, suggesting increased developmental or environmental sensitivity of CHH methylation. Genome-wide methylation levels calculated for each context showed a general increase from Spring to Summer and Autumn (Figure 1c). Offspring samples also displayed higher methylation levels than Spring samples across all three contexts (Figure 1c) with a genome-wide level of methylation more like the Summer and Autumn conditions.

**Figure 1.**
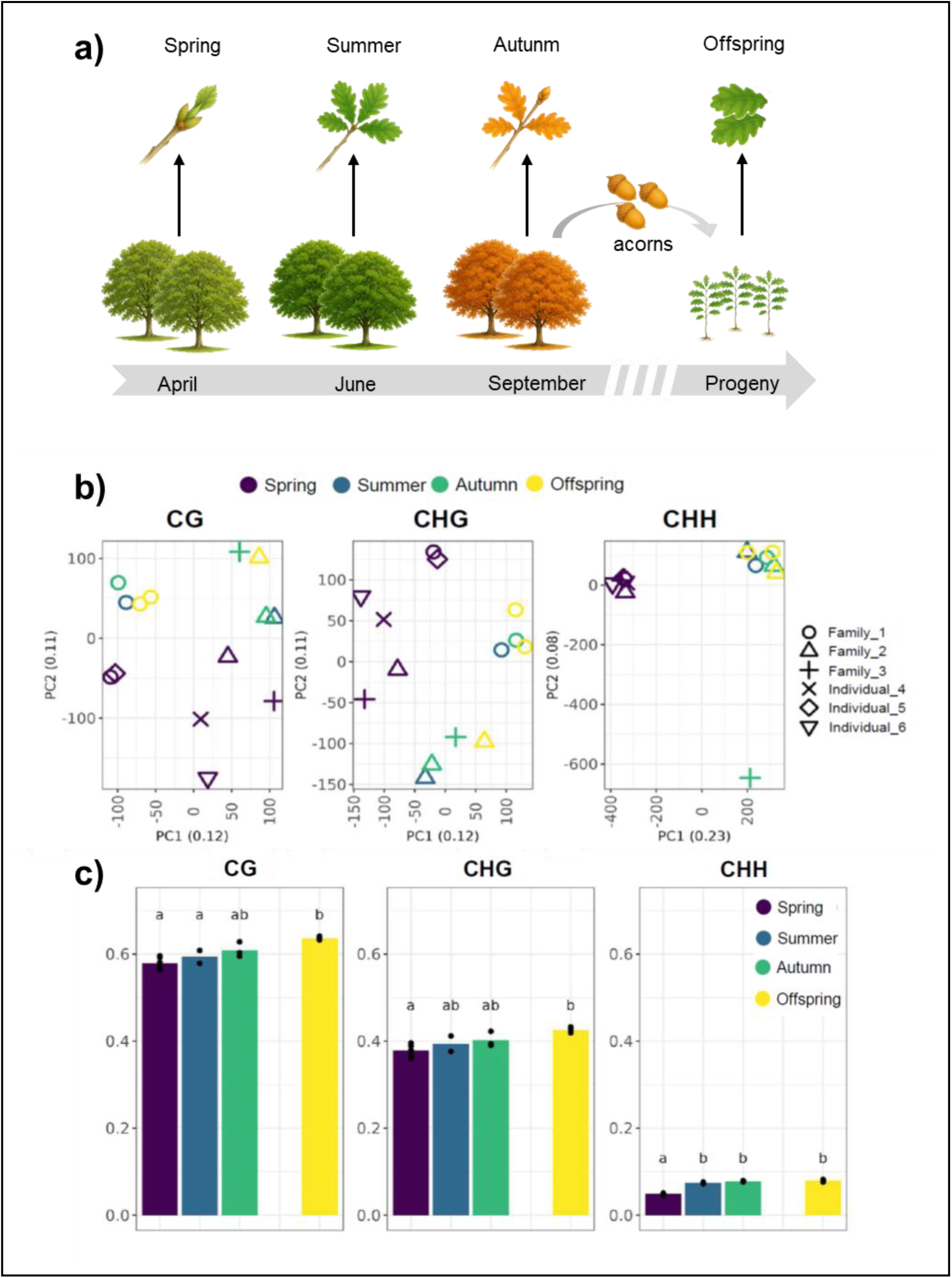
Seasonal and generational dynamics of DNA methylation in *Quercus robur*. (**a**) Scheme representing the samples collected for this analysis. (**b**) Principal component analysis (PCA) of DNA methylation profiles across CG, CHG, and CHH contexts, respectively. (**c**) Barplots showing genome-wide methylation levels across CG, CHG, and CHH contexts, respectively. Black circles indicate single replicates (i.e. each individual tree). Different letters indicate statistically significant differences between groups (Pair-wise T-Test; p < 0.05; n = 2-6).

Chromosome-specific analyses mirrored the genome-wide trends (Figures S3a-b). In all contexts, methylation levels followed the same broad pattern throughout the genome. However, methylation was lowest in Spring and increased toward Summer and Autumn. Again, CHH methylation showed the most notable seasonal dynamics, with a clear increase from Spring to Autumn, indicating that the seasonal effect is extended to the entire genome.

To further analyse the relation of epigenetic and genetic variation, structural variants (SNPs) were called directly from the WGBS datasets and used to perform PCA (Figure S2b). We observed that the samples from tree id 6387 (Family 1) are more differentiated than the ids 8749 and 8673 (Family 2 and 3, respectively), and on this clustering the grouping of samples by Spring (Figure 1b) was lost. This suggest that the effect we observed for DNA methylation does not reflect genetic variation occurring across the season, or between different tree branches. Indeed, this SNPs-based clustering is highly similar to the DArTseq analysis (Figure S2a), confirming that the seasonal effects observed (Figure 1b-c) is due to pure epigenetic variation and do not correlate with genetic differences.

### Seasonal Shifts in Oak DNA Methylation Are Dominated by CHH Dynamics

To investigate the variation of DNA methylation observed across seasons, we called differentially methylated regions (DMRs) among all pairwise Season combinations for each of the three cytosine contexts (CG, CHG and CHH). For all contexts, we identified a higher number of Seasonal DMRs in the comparisons involving Spring, and lower numbers when Summer and Autumn were compared (Figures 2a and S4a,b). In the comparison involving Spring we observed a higher proportion of CG and CHG DMRs losing methylation than the DMRs from the comparison of other seasons (Figure S4a,b, respectively). Virtually all Spring DMRs in the CHH context are gaining methylation (Figure 2a) in Summer (100%) and Autumn (99.9%), consistent with the genome-wide increase of CHH methylation from Spring to other seasons (Figure S4).

**Figure 2.**
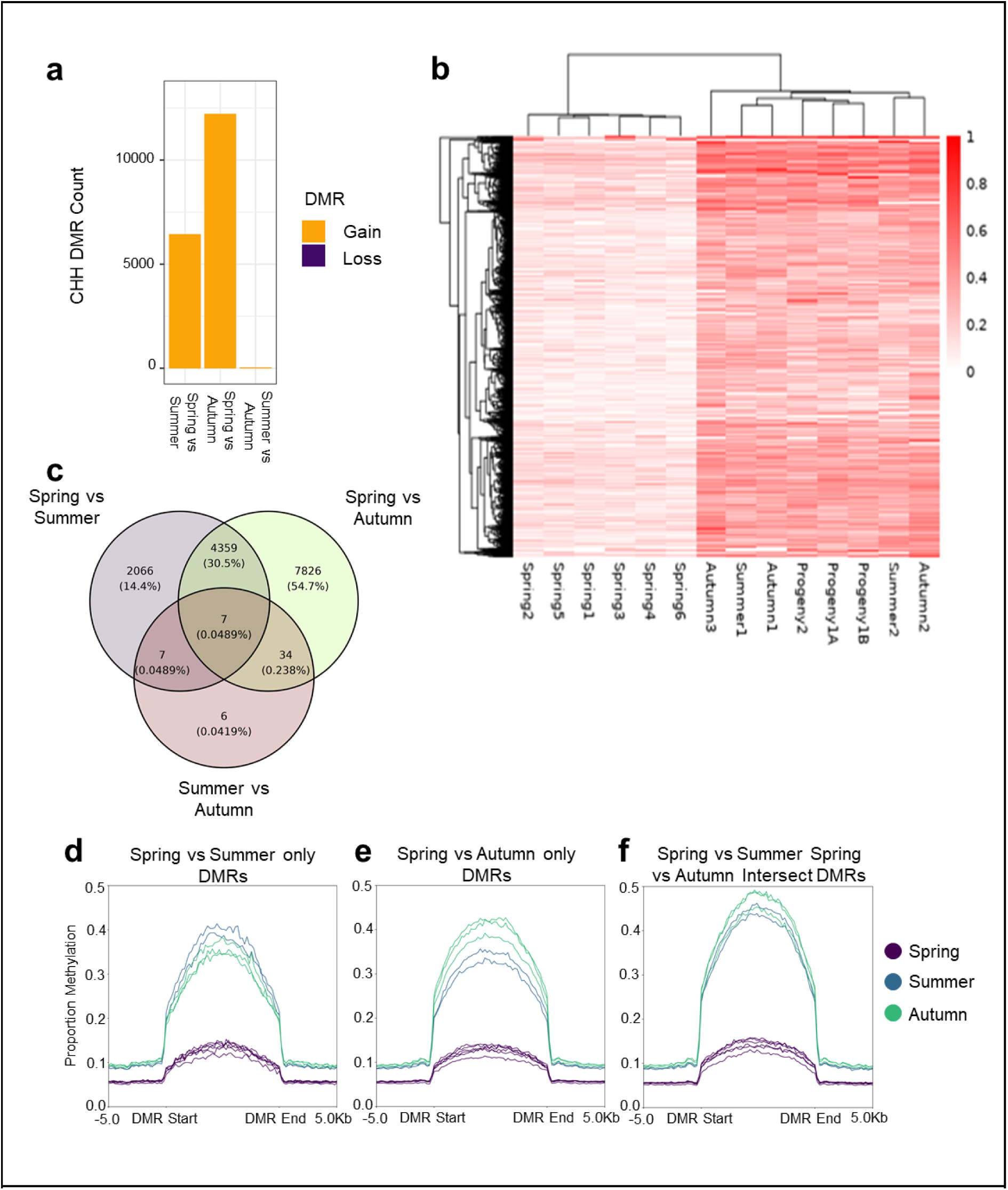
CHH methylation dynamic increases across seasons. **(a)** Bar plots displaying the number of CHH Differentially Methylated Region (DMRs) identified in seasonal comparisons. **(b)** Heatmap displaying the CHH methylation values in DMRs across each individual sample. **(c)** Venn diagram displaying the degree of overlap among the number of DMRs called in each seasonal comparison. The proportion of DMR in each area displayed in parentheses as a percentage of all DMRs. **(d-f)** Cumulative CHH methylation calculated by aligning Spring CHH DMRs, in each seasonal sample, separated by regions found significantly differentially methylated only in the Spring versus (vs) Summer comparison **(d)**, only in the Spring vs Autumn comparison **(e)**, or common to both comparisons **(f)**.

We computed methylation levels for all samples at loci where Seasonal DMRs were identified in all cytosine contexts and clustered by sample and locus. In the CG and CHG contexts, the samples were grouped by genotype and offspring (Figure S5), suggesting that the methylation patterns may be associated with the sample genotype. In contrast, in the CHH context, there was a clear clustering of Spring samples together, which are characterised by a marked hypomethylation (Figure 2b) when compared to Summer and Autumn.

To account for genetic and epigenetic variation among trees, we identified “genetic DMRs” by comparing samples from the same tree across seasons (Figure S6). These genetic DMRs showed minimal overlap with seasonal DMRs: ∼0.2%, ∼0.4%, and ∼0.14% for CG, CHG and CHH contexts, respectively (Figure S6a-c), indicating the family-level clustering observed for CG and CHG contexts (Figure S6d,e) likely reflecting genotype-specific seasonal responses rather than direct genetic differences.

Seasonal DMRs showed substantial overlap across all sequence contexts, particularly for CHH DMRs between Spring vs Summer and Spring vs Autumn comparisons (30.5%; Figure 2c, Figure S6d,e). These loci generally gained methylation from Spring to the later seasons. Averaged CHH methylation profiles (Figure 2d-f) confirmed consistently low methylation in Spring and higher levels thereafter, indicating that CHH DMRs mark genomic regions with increasing methylation over the seasonal cycle, most prominently from Spring to Summer.

We also noticed that, contrary to CHH DMRs, several CG DMRs showed decreased methylation over the season (Figure S4a), leading us to analyse CHH methylation at the same loci to investigate a relation between the two epigenetic marks. Indeed, CG methylation in seasonal CHH DMRs was consistently higher in spring (Figure S7). Among the most methylated seasonal CG DMRs (>0.4 relative methylation), a spring-specific clade was evident (Figure S8a), while CHH methylation in these regions showed an inverse pattern, maintaining spring sample clustering (Figure S8b). These results indicate seasonal interplay between epigenetic pathways, with CHH hypermethylation potentially compensating for CG hypomethylation from spring to summer/autumn.

### Seasonal CHH Hypermethylation Is Linked to Promoter Regions and TIR Elements

To investigate potential functional roles of DNA hypermethylation over seasons, the overlap of CHH Spring DMRs with oak genetic features was performed (Figure 3a). We found that CHH gain of methylation over the Summer and Autumn occurs mostly on gene promoters with a decrease methylation in gene bodies, while variation in methylation in CG and CHG contexts do not show substantial trends (Figure S9). Genes with promoters overlapping Seasonal CHH DMRs were enriched for Gene Ontology (GO) categories related to leaf development, photomorphogenesis, and hormonal signalling (Figure 3b), suggesting that DNA methylation may help regulate genes involved in environmental responses and leaf development.

**Figure 3.**
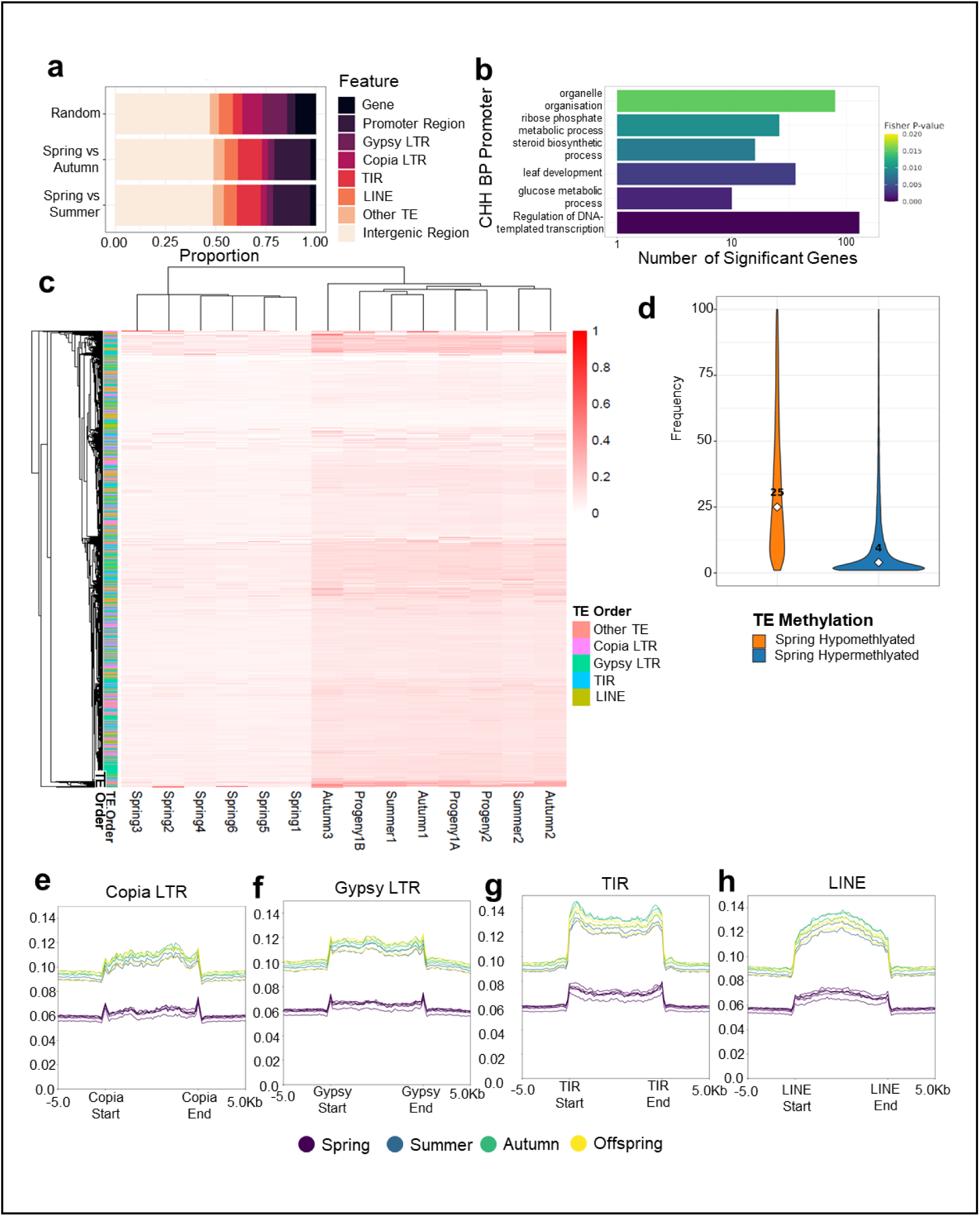
Seasonal CHH Differentially Methylation Regions (DMRs) are associated with gene promoters and Transposable Elements (TEs). **(a)** Overlaps of genetic features and Spring DMRs in the CHH. **(b)** Gene ontology from promoters at CHH context. **(c)** Clustering of methylation in TEs in the CHH. **(d)** The 10% most hyper and hypo methylated TEs in Spring were identified and their frequency computed. (**e-h**) Methylation of Copia TEs, Gypsy TEs, LINE TEs and TIR TEs in all seasons.

Seasonal CHH DMRs were predominantly enriched in TIR TEs, with fewer overlaps in LTR TEs (Figure 3a). CHH methylation analysis showed Spring samples forming a distinct cluster with reduced methylation, without clear grouping by TE type (Figure 3c). In oak, CHH methylation of common TEs (Copia, Gypsy, TIR, and LINE elements) increased in Summer and Autumn (Figure 3e–h), and TIR TEs were generally closer to genes, particularly those overlapping DMRs (Figures S10, S11), suggesting a potential role in gene regulation. Comparing the 10% most and least hypermethylated TEs revealed that those with the largest CHH changes were also more frequently repeated in the genome (Figure 3d), indicating that genome repetitiveness may facilitate seasonal epigenetic variation.

### Offspring Differentially Methylation Regions (DMRs) is Associated with Promoters and Gene Bodies in the CG and CHG Contexts

We then examined epigenetic differences between Offspring and parental trees to identify Generational DMRs. To ensure anatomical and molecular consistency and avoid artefacts from high methylation variation in Spring samples, we excluded this season from the analysis. Unlike previously observed seasonal methylation patterns, the Generational DMRs mainly showed changes in the CG and CHG contexts (Figure 4a). Both CG and CHG generational DMRs were overlapped with genetic features (Figure 4b). In the CHG context, we found an overrepresentation of Intergenic Regions in both the gain and loss types and an underrepresentation in the Genes (Fisher’s exact test; p < 0.001). In the CG context, there was an over-representation of both Intergenic Regions and Genes. In all contexts, we observed underrepresentation of LTR TEs, such as Copia TEs and Gypsy TEs.

**Figure 4.**
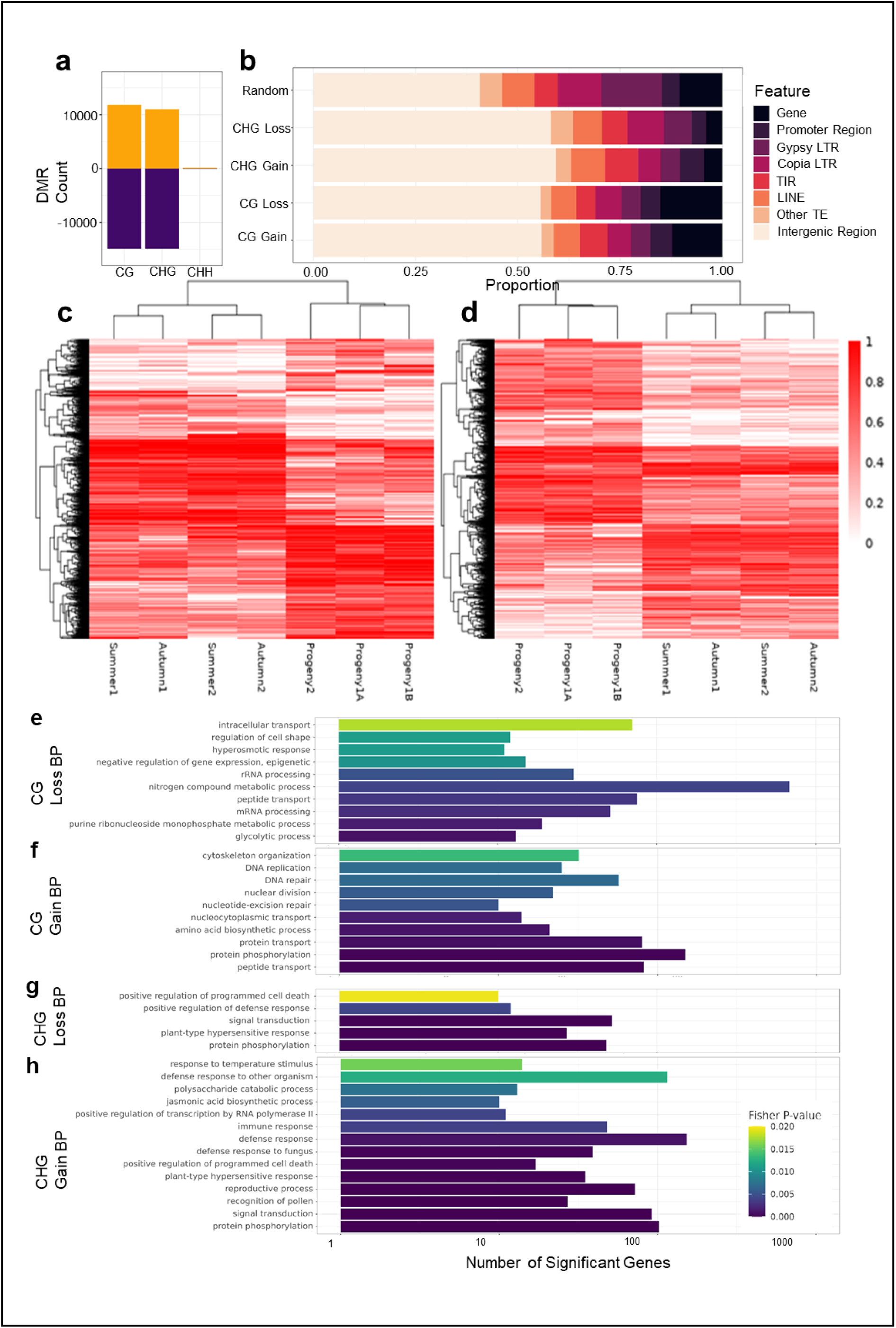
Generational CG and CHG Differentially Methylated Regions (DMRs) are associated with defence and stress responses. **(a)** DMRs were called comparing Offspring and their parents in Summer and Autumn in the CG, CHG and CHH contexts. **(b)** DMRs overlapping with genetic elements were identified with overrepresentation of intergenic regions in CG and CHG genes in the CG context and underrepresentation of the Gypsy TEs and Copia TEs elements. Heatmaps of the DMRs in the CG **(c)** and CHG **(d).** Ontologies representing the Biological Processes (BP) of the genes overlapping with CG (hypomethylation **(e)** and hypermethylation **(f)**) and CHG (hypomethylation **(g)** and hypermethylation **(h)**) DMRs.

A hierarchical clustering analysis of the generational DMR loci highlights a clear differential grouping of Offspring and parent samples, suggesting that DMRs identified in this analysis are indeed associated with generational effects (Figure 4c, d). To reveal the genes overlapping with Generational DMRs, we identified ontologies of genes overlapping both CG and CHG contexts DMRs for both gain and loss in methylation (Figure 4e-h). We observed a clear overrepresentation of ontologies associated with defence responses as well as reproductive regulation, which could reflect a combination of developmental differences associated with age, as well as the different environmental conditions of parental trees living in the wild compared to their offspring, which have been grown in controlled conditions.

## Discussion

This study provides the first genome-wide analysis of seasonal and generational DNA methylation dynamics in *Quercus robur*, a keystone of European temperate forests. By combining temporal sampling of mature trees with comparisons to their offspring, we reveal that the oak methylome is both environmentally dynamic and developmentally programmed. Two major results emerged: (i) CHH methylation displays strong seasonal plasticity linked to phenological transitions, and (ii) CG and CHG methylation mark generational reprogramming in seedlings, particularly in genes related to development and defence. These variations appear to be largely independent by genetic variation in trees, which has been recently link to a certain degree of variation in DNA methylation in long living trees (Zhou *et al*., 2024; Johannes, 2025)

The strongest seasonal shifts occurred in the CHH context, with methylation consistently lower in Spring and increasing through Summer and Autumn. This pattern reflects the dynamic nature of CHH methylation observed in other plants (Zhang *et al*., 2006; Bouyer *et al*., 2017) but here is captured in a long-lived tree with a deciduous growth cycle. The Spring-Summer rise coincides with bud burst and leaf expansion, suggesting that CHH hypermethylation contributes to transcriptional reprogramming as leaves transition from dormant to fully functional photosynthetic organs. Conversely, the stabilization of methylation in Autumn may relate to preparation for leaf senescence and nutrient remobilisation.

Seasonal CHH changes were often accompanied by a decrease in CG methylation, revealing an inverse relationship between contexts. This reciprocal regulation, previously reported in Arabidopsis (Borges *et al*., 2012) and in tomato and soybean (Zhang *et al*., 2018; Nyikó *et al*., 2025), may reflect compensatory interactions among epigenetic pathways that coordinate developmental stage transitions. In oaks, such a mechanism could provide flexibility to adjust growth and metabolic investment according to environmental cues while maintaining overall genomic stability.

The non-random genomic distribution of seasonal differentially methylated regions (DMRs) suggests that these loci hold regulatory importance. In *Arabidopsis halleri*, methylome profiling across natural seasonal cycles showed that cytosine methylation patterns remain largely stable but exhibit context-specific fluctuations at defined genomic regions, particularly near transcriptional start sites of environmentally responsive (Ito *et al*., 2019). Similarly, in *Populus trichocarpa*, genome-wide methylation analyses have revealed that DMRs are enriched in promoter regions and correlate with transcriptional variation across diel and environmental conditions, highlighting a potential role for methylation in transcriptional tuning (Liang *et al*., 2019). Functional enrichment analyses of methylation-associated genes point to processes such as organelle organisation, carbon metabolism, and morphogenesis, core components of phenological adaptation and leaf developmental transitions. Collectively, these findings suggest that environmentally sensitive DNA methylation may serve as an epigenetic interface aligning transcriptional activity with seasonal metabolic and structural demands.

The preferential targeting of TIR transposable elements, rather than LTR retrotransposons, provides an additional layer of regulation. TIRs are frequently positioned near genes and have been implicated in environmentally induced chromatin remodelling (Iwasaki & Paszkowski, 2014; Catoni *et al*., 2018a; Emmerson & Catoni, 2025; Mencia *et al*., 2025; Tossolini *et al*., 2025). Their association with seasonal CHH methylation in oak suggests they may act as epigenetic switches, modulating the chromatin environment around stress- or development-related genes. In this way, TEs could serve not only as targets for silencing but also as facilitators of rapid regulatory responses, an especially valuable feature for perennial trees facing annual cycles of environmental change.

In contrast to seasonal plasticity in mature trees, progeny displayed distinct methylation profiles dominated by CG and CHG contexts. These generational DMRs were enriched in genes, particularly those linked to development and defence responses. The clustering of progeny separately from both their parents and seasonal samples indicates that substantial reprogramming occurs during the transition from seed to seedling. This is consistent with reports in other long-lived plants where maternal inputs and local conditions shape early-life epigenetic states (Kakutani, 2002; Iwasaki & Paszkowski, 2014; Kawakatsu *et al*., 2017).

Functionally, contrasts in methylation profiles between seedlings and mature trees are likely driven by environmental rather than ontogenetic factors. Seedlings grown under controlled, low-stress conditions experience minimal biotic challenge, whereas field-grown trees are continually exposed to microbial and herbivory pressures that can reshape methylation landscapes (Dowen *et al*., 2012; Secco *et al*., 2015). In oaks and other long-lived perennials, DNA methylation has been shown to respond dynamically to localised stress, particularly within CHG contexts associated with defence-related genes (Gugger *et al*., 2013; Liang *et al*., 2019). The enrichment of CG methylation among genes involved in primary metabolism and RNA processing instead suggests a conserved role in maintaining cellular homeostasis during active growth (Zhang *et al*., 2018). Collectively, these patterns support the view that environmental context exerts a dominant influence on the distribution of methylation variants, cautioning against the interpretation of defence-related epigenetic differences as age-dependent traits.

Taken together, our findings suggest that oaks deploy a multi-layered epigenetic strategy shaped by their perennial life history. Seasonal CHH methylation provides flexibility, allowing adult trees to recalibrate gene expression each year in response to environmental cues. Generational differences in CG and CHG methylation, on the other hand, establish developmental priorities suited to early life, particularly defence investment. This combination of plasticity and stability may help reconcile the dichotomy between herbaceous plants, which must adapt to immediate conditions, and long-lived trees, which must persist for centuries under shifting climates (Zhang *et al*., 2018).

Such dynamics also raise the question of epigenetic memory. Long-lived species may accumulate methylation patterns reflecting prior environments, which could influence future phenological responses. At the same time, the reprogramming observed in progeny suggests that not all marks are transmitted across generations. Distinguishing between reversible, plastic modifications and stable, heritable marks remains an essential challenge in understanding the adaptive potential of tree epigenomes.

Our study is limited to leaf tissue, which provides a window into seasonal growth but may not capture methylation changes in reproductive or meristematic tissues where transgenerational inheritance is most likely to occur (Johannes, 2025). In addition, while our sampling controlled for genetic uniformity, we could not control paternal contributions to progeny epigenomes. Future studies using controlled crosses and reciprocal transplant experiments will be critical for disentangling genetic and environmental influences.

Our results indicate that *Quercus robur* exhibits DNA methylation variation both across seasons and between generations, with CHH methylation fluctuating seasonally and CG/CHG methylation differing between seedlings and adult trees. These context-dependent methylation dynamics underscore the importance of environmental and developmental factors in shaping the epigenome of long-lived perennials. Looking forward, such methylation variation may represent a key component of forest tree adaptation, buffering populations against short-term environmental variability while influencing developmental trajectories across generations. As climate change intensifies, understanding the interplay between epigenetic plasticity and inheritance will be critical for predicting and managing the resilience of forest ecosystems.

## Data Availability Statement

Sequencing data, list of DMRs generated and the CX reports files have been deposited in Gene Expression Omnibus (https://www.ncbi.nlm.nih.gov/geo/) under the accession number GSE303740. The computations described in this paper were performed using the University of Birmingham’s BlueBEAR HPC service, which provides the High Performance Computing service to the University’s research community. See http://www.birmingham.ac.uk/bear for more details. The full code developed is available on Github (https://github.com/PlantPriming/Oak_Seasons).

## Author Contribution

RSL, EL and MCa designed research; RLS and JH performed research; RLS, MCa, JH, MCh, JB, AABL and KSO analysed data; EL and MCa contributed with reagents/analytic tools; KH and ARM managed experimental site and provided technical support; EL and MCa supervised research; EL and MCa acquired funding; RSL, JH, EL, MCa, wrote the paper; all authors contributed to the final manuscript.

## Acknowledgments

This work was supported by the UK Research and Innovation (UKRI) Future of UK Treescapes programme through the MEMBRA project (grant number NE/V021346/1). We are extremely grateful to the Birmingham Institute of Forest Research (BIFoR) for providing access to infrastructure and technical support. We also thank the arborists and technical staff at BIFoR FACE for their assistance with sample collection and tree monitoring.

## References

Amasino R. 2010. Seasonal and developmental timing of flowering. The Plant Journal 61(6): 1001–1013.

Andrés F, Coupland G. 2012. The genetic basis of flowering responses to seasonal cues. Nature Reviews Genetics 13(9): 627–639.

Ashley MV. 2021. Answers Blowing in the Wind: A Quarter Century of Genetic Studies of Pollination in Oaks. Forests 12(5): 575.

Bartee L, Malagnac F, Bender J. 2001. Arabidopsis cmt3 chromomethylase mutations block non-CG methylation and silencing of an endogenous gene. Genes Dev 15(14): 1753–1758.

Benito Garzón M, Baillou F, Costa e Silva F, Faria C, Marchi M, Vendramin GG, Vizcaíno-Palomar N. 2024. Warmer springs favour early germination of range-wide Quercus suber L. populations. European Journal of Forest Research 143(1): 157–168.

Bölöni J, Aszalós R, Tamás F, Ódor P. 2021. Forest type matters: Global review about the structure of oak dominated old-growth temperate forests. Forest Ecology and Management 500: 119629.

Borges F, Calarco JP, Martienssen RA. 2012. Reprogramming the epigenome in Arabidopsis pollen. Cold Spring Harb Symp Quant Biol 77: 1–5.

Bouyer D, Kramdi A, Kassam M, Heese M, Schnittger A, Roudier F, Colot V. 2017. DNA methylation dynamics during early plant life. Genome Biology 18(1): 179.

Bratu I, Dinca L, Constandache C, Murariu G. 2025. Resilience and Decline: The Impact of Climatic Variability on Temperate Oak Forests. Climate 13(6): 119.

Browne L, Wright JW, Fitz-Gibbon S, Gugger PF, Sork VL. 2019. Adaptational lag to temperature in valley oak (*Quercus lobata*) can be mitigated by genome-informed assisted gene flow. Proceedings of the National Academy of Sciences 116(50): 25179–25185.

Burnett AC, Serbin SP, Lamour J, Anderson J, Davidson KJ, Yang D, Rogers A. 2021. Seasonal trends in photosynthesis and leaf traits in scarlet oak. Tree Physiology 41(8): 1413–1424.

Cao X, Jacobsen SE. 2002. Role of the arabidopsis DRM methyltransferases in de novo DNA methylation and gene silencing. Curr Biol 12(13): 1138–1144.

Catoni M, Griffiths J, Becker C, Zabet NR, Bayon C, Dapp M, Lieberman-Lazarovich M, Weigel D, Paszkowski J. 2017. DNA sequence properties that predict susceptibility to epiallelic switching. The EMBO Journal 36(5): 617–628.

Catoni M, Jonesman T, Cerruti E, Paszkowski J. 2018a. Mobilization of Pack-CACTA transposons in Arabidopsis suggests the mechanism of gene shuffling. Nucleic Acids Research 47(3): 1311–1320.

Catoni M, Tsang JM, Greco AP, Zabet Nicolae R. 2018b. DMRcaller: a versatile R/Bioconductor package for detection and visualization of differentially methylated regions in CpG and non-CpG contexts. Nucleic Acids Research 46(19): e114–e114.

Catoni M, Zabet NR. 2021. Analysis of Plant DNA Methylation Profiles Using R. Methods Mol Biol 2250: 219–238.

Chen S, Zhou Y, Chen Y, Gu J. 2018. fastp: an ultra-fast all-in-one FASTQ preprocessor. Bioinformatics 34(17): i884–i890.

Conrad AO, McPherson BA, Lopez-Nicora HD, D’Amico KM, Wood DL, Bonello P. 2019. Disease incidence and spatial distribution of host resistance in a coast live oak/sudden oak death pathosystem. Forest Ecology and Management 433: 618–624.

Dowen RH, Pelizzola M, Schmitz RJ, Lister R, Dowen JM, Nery JR, Dixon JE, Ecker JR. 2012. Widespread dynamic DNA methylation in response to biotic stress. Proceedings of the National Academy of Sciences 109(32): E2183–E2191.

Egea LA, Mérida-García R, Kilian A, Hernandez P, Dorado G. 2017. Assessment of Genetic Diversity and Structure of Large Garlic (Allium sativum) Germplasm Bank, by Diversity Arrays Technology “Genotyping-by-Sequencing” Platform (DArTseq). Frontiers in Genetics Volume 8–2017.

Emmerson R, Catoni M. 2025. The role of mobile DNA elements in the dynamics of plant genome plasticity. Journal of Experimental Botany 76(9): 2433–2446.

Gugger PF, Ikegami M, Sork VL. 2013. Influence of late Quaternary climate change on present patterns of genetic variation in valley oak, Quercus lobata Née. Molecular Ecology 22(13): 3598–3612.

Guo W, Zhu P, Pellegrini M, Zhang MQ, Wang X, Ni Z. 2017. CGmapTools improves the precision of heterozygous SNV calls and supports allele-specific methylation detection and visualization in bisulfite-sequencing data. Bioinformatics 34(3): 381–387.

Han B, Wang L, Xian Y, Xie X-M, Li W-Q, Zhao Y, Zhang R-G, Qin X, Li D-Z, Jia K-H. 2022. A chromosome-level genome assembly of the Chinese cork oak (Quercus variabilis). Frontiers in Plant Science Volume 13–2022.

Haneca KaB, Hans,. 2009. Oaks, tree-rings and wooden cultural heritage: a review of the main characteristics and applications of oak dendrochronology in Europe. Journal of Archaeological Science 36(1): 1–11.

Hart KM, Curioni G, Blaen P, Harper NJ, Miles P, Lewin KF, Nagy J, Bannister EJ, Cai XM, Thomas RM, et al. 2020. Characteristics of free air carbon dioxide enrichment of a northern temperate mature forest. Global Change Biology 26(2): 1023–1037.

Ito T, Nishio H, Tarutani Y, Emura N, Honjo MN, Toyoda A, Fujiyama A, Kakutani T, Kudoh H. 2019. Seasonal Stability and Dynamics of DNA Methylation in Plants in a Natural Environment. Genes 10(7): 544.

Iwasaki M, Paszkowski J. 2014. Identification of genes preventing transgenerational transmission of stress-induced epigenetic states. Proceedings of the National Academy of Sciences 111(23): 8547–8552.

Jato V, Rodríguez-Rajo FJ, Fernandez-González M, Aira MJ. 2015. Assessment of Quercus flowering trends in NW Spain. International Journal of Biometeorology 59(5): 517–531.

Johannes F. 2025. Branching architecture limits the number of fixed somatic mutations in trees. G3 Genes|Genomes|Genetics.

Kakutani T. 2002. Epi-Alleles in Plants: Inheritance of Epigenetic Information over Generations. Plant and Cell Physiology 43(10): 1106–1111.

Kapoor B, Jenkins J, Schmutz J, Zhebentyayeva T, Kuelheim C, Coggeshall M, Heim C, Lasky JR, Leites L, Islam-Faridi N, et al. 2023. A haplotype-resolved chromosome-scale genome for Quercus rubra L. provides insights into the genetics of adaptive traits for red oak species. G3 Genes|Genomes|Genetics 13(11).

Kawakatsu T, Nery JR, Castanon R, Ecker JR. 2017. Dynamic DNA methylation reconfiguration during seed development and germination. Genome Biology 18(1): 171.

Kilian A, Wenzl P, Huttner E, Carling J, Xia L, Blois H, Caig V, Heller-Uszynska K, Jaccoud D, Hopper C, et al. 2012. Diversity Arrays Technology: A Generic Genome Profiling Technology on Open Platforms. In: Pompanon F, Bonin A eds. Data Production and Analysis in Population Genomics: Methods and Protocols. Totowa, NJ: Humana Press, 67–89.

Krueger F, Andrews SR. 2011. Bismark: a flexible aligner and methylation caller for Bisulfite-Seq applications. Bioinformatics 27(11): 1571–1572.

Labella-Ortega M, Martín C, Valledor L, Castiglione S, Castillejo M-Á, Jorrín-Novo JV, Rey M-D. 2024. Unravelling DNA methylation dynamics during developmental stages in Quercus ilex subsp. ballota [Desf.] Samp. BMC Plant Biology 24(1): 823.

Langmead B, Salzberg SL. 2012. Fast gapped-read alignment with Bowtie 2. Nature Methods 9(4): 357–359.

Lawrence M, Huber W, Pagès H, Aboyoun P, Carlson M, Gentleman R, Morgan MT, Carey VJ. 2013. Software for computing and annotating genomic ranges. PLoS Comput Biol 9(8): e1003118.

Liang L, Chang Y, Lu J, Wu X, Liu Q, Zhang W, Su X, Zhang B. 2019. Global Methylomic and Transcriptomic Analyses Reveal the Broad Participation of DNA Methylation in Daily Gene Expression Regulation of Populus trichocarpa. Front Plant Sci 10: 243.

López-Hidalgo C, Guerrero-Sánchez VM, Gómez-Gálvez I, Sánchez-Lucas R, Castillejo-Sánchez MA, Maldonado-Alconada AM, Valledor L, Jorrín-Novo JV. 2018. A Multi-Omics Analysis Pipeline for the Metabolic Pathway Reconstruction in the Orphan Species Quercus ilex. Frontiers in Plant Science Volume 9–2018.

Madritsch S, Wischnitzki E, Kotrade P, Ashoub A, Burg A, Fluch S, Brüggemann W, Sehr EM. 2019. Elucidating Drought Stress Tolerance in European Oaks Through Cross-Species Transcriptomics. G3 Genes|Genomes|Genetics 9(10): 3181–3199.

Mencia R, Arce AL, Houriet C, Xian W, Contreras A, Shirsekar G, Weigel D, Manavella PA. 2025. Transposon-triggered epigenetic chromatin dynamics modulate EFR-related pathogen response. Nature Structural & Molecular Biology 32(1): 199–211.

Miguel A, de Vega-Bartol J, Marum L, Chaves I, Santo T, Leitão J, Varela MC, Miguel CM. 2015. Characterization of the cork oak transcriptome dynamics during acorn development. BMC Plant Biology 15(1): 158.

Nyikó T, Gyula P, Ráth S, Sós-Hegedűs A, Csorba T, Abbas SH, Bóka K, Pettkó-Szandtner A, Móricz ÁM, Molnár BP, et al. 2025. INCREASED DNA METHYLATION 3 forms a potential chromatin remodelling complex with HAIRPLUS to regulate DNA methylation and trichome development in tomato. The Plant Journal 121(6): e70085.

Peck LD, Sork VL. 2024. Can DNA methylation shape climate response in trees? Trends in Plant Science 29(10): 1089–1102.

Plomion C, Aury J-M, Amselem J, Leroy T, Murat F, Duplessis S, Faye S, Francillonne N, Labadie K, Le Provost G, et al. 2018. Oak genome reveals facets of long lifespan. Nature Plants 4(7): 440–452.

Pockrandt C, Alzamel M, Iliopoulos CS, Reinert K. 2020. GenMap: ultra-fast computation of genome mappability. Bioinformatics 36(12): 3687–3692.

Purcell S, Neale B, Todd-Brown K, Thomas L, Ferreira MA, Bender D, Maller J, Sklar P, de Bakker PI, Daly MJ, et al. 2007. PLINK: a tool set for whole-genome association and population-based linkage analyses. Am J Hum Genet 81(3): 559–575.

Ramírez F, Dündar F, Diehl S, Grüning BA, Manke T. 2014. deepTools: a flexible platform for exploring deep-sequencing data. Nucleic Acids Research 42(W1): W187–W191.

Ramos AM, Usié A, Barbosa P, Barros PM, Capote T, Chaves I, Simões F, Abreu I, Carrasquinho I, Faro C, et al. 2018. The draft genome sequence of cork oak. Scientific Data 5(1): 180069.

Rey M-D, Labella-Ortega M, Guerrero-Sánchez VM, Carleial R, Castillejo MÁ, Ruggieri V, Jorrín-Novo JV. 2023. A first draft genome of holm oak (Quercus ilex subsp. ballota), the most representative species of the Mediterranean forest and the Spanish agrosylvopastoral ecosystem “dehesa”. Frontiers in Molecular Biosciences Volume 10–2023.

Rudy E, Tanwar UK, Szlachtowska Z, Grabsztunowicz M, Arasimowicz-Jelonek M, Sobieszczuk-Nowicka E. 2024. Unveiling the role of epigenetics in leaf senescence: a comparative study to identify different epigenetic regulations of senescence types in barley leaves. BMC Plant Biology 24(1): 863.

Sanchez-Lucas R, Bosanquet JL, Henderson J, Catoni M, Pastor V, Luna E. 2025. Elicitor Specific Mechanisms of Defence Priming in Oak Seedlings Against Powdery Mildew. Plant, Cell & Environment 48(6): 4455–4474.

Sanchez-Lucas R, Mayoral C, Raw M, Mousouraki MA, Luna E. 2023. Elevated CO2 alters photosynthesis, growth and susceptibility to powdery mildew of oak seedlings. Biochem J 480(17): 1429–1443.

Secco D, Wang C, Shou H, Schultz MD, Chiarenza S, Nussaume L, Ecker JR, Whelan J, Lister R. 2015. Stress induced gene expression drives transient DNA methylation changes at adjacent repetitive elements. eLife 4: e09343.

Silva HG, Sobral R, Alhinho AT, Afonso HR, Ribeiro T, Silva PMA, Bousbaa H, Morais-Cecílio L, Costa MMR. 2024. Genetic and epigenetic control of dormancy transitions throughout the year in the monoecious cork oak. Physiologia Plantarum 176(6): e14620.

Song Q, Chen ZJ. 2015. Epigenetic and developmental regulation in plant polyploids. Current Opinion in Plant Biology 24: 101–109.

Sork VL, Fitz-Gibbon ST, Puiu D, Crepeau M, Gugger PF, Sherman R, Stevens K, Langley CH, Pellegrini M, Salzberg SL. 2016. First Draft Assembly and Annotation of the Genome of a California Endemic Oak Quercus lobata Née (Fagaceae). G3 Genes|Genomes|Genetics 6(11): 3485–3495.

Tossolini I, Mencia R, Arce AL, Manavella PA. 2025. The genome awakens: transposon-mediated gene regulation. Trends in Plant Science 30(8): 857–871.

Zhang H, Lang Z, Zhu J-K. 2018. Dynamics and function of DNA methylation in plants. Nature Reviews Molecular Cell Biology 19(8): 489–506.

Zhang X, Yazaki J, Sundaresan A, Cokus S, Chan SW, Chen H, Henderson IR, Shinn P, Pellegrini M, Jacobsen SE, et al. 2006. Genome-wide high-resolution mapping and functional analysis of DNA methylation in arabidopsis. Cell 126(6): 1189–1201.

Zheng X, Levine D, Shen J, Gogarten SM, Laurie C, Weir BS. 2012. A high-performance computing toolset for relatedness and principal component analysis of SNP data. Bioinformatics 28(24): 3326–3328.

Zhou M, Schmied G, Bradatsch M, Resente G, Hazarika R, Kakoulidou I, Costa M, Serra M, Uhl E, Schmitz R, et al. 2024. Accelerated growth increases the somatic epimutation rate in trees. bioRxiv: 2024.2005.2007.592680.

Zilberman D, Gehring M, Tran RK, Ballinger T, Henikoff S. 2007. Genome-wide analysis of Arabidopsis thaliana DNA methylation uncovers an interdependence between methylation and transcription. Nature Genetics 39(1): 61–69.

