## Supplementary Figures for "Epigenomic Landscape of Oak (*Quercus robur*) across Seasons and Generations"

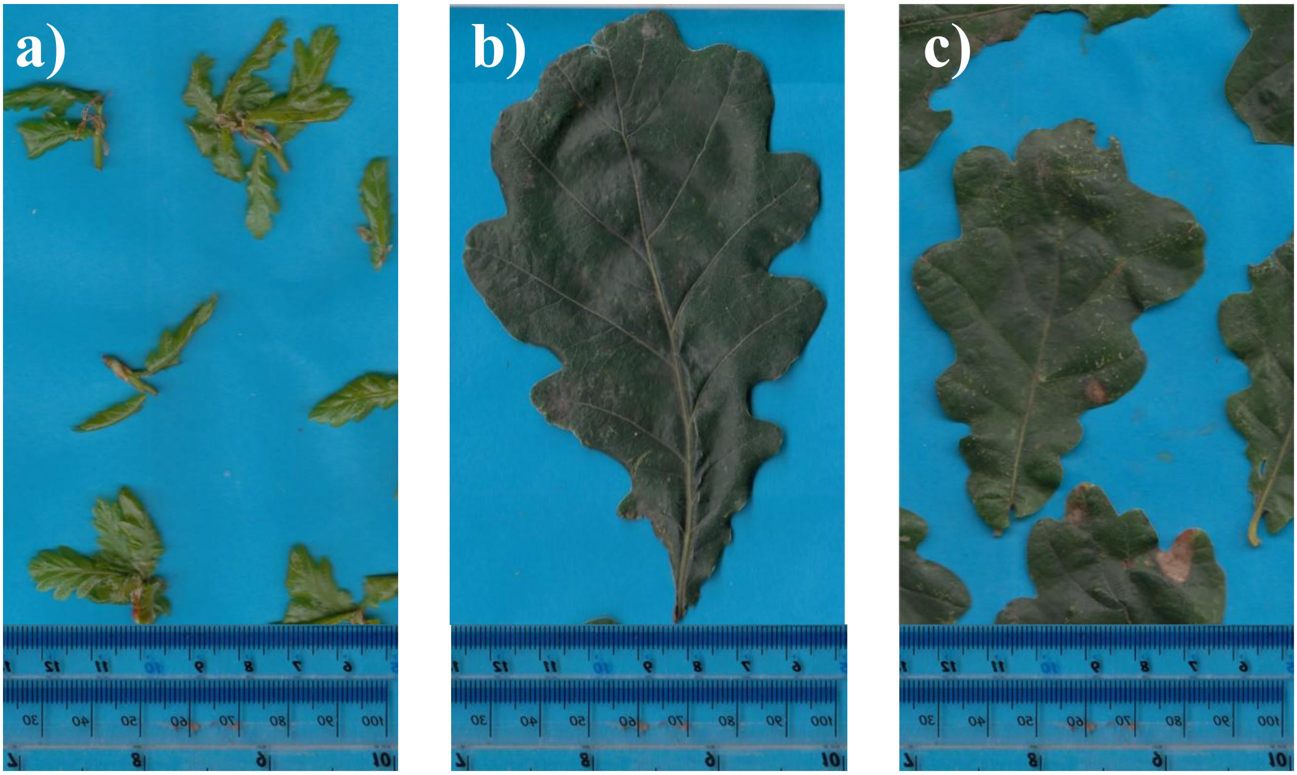


**Figure Supplementary 1**. Representative leaves from the sampling collection in 2021 corresponding to **a)** April, **b)** June and **c)** September. All pictures have been performed at the same scale and imaging settings by using a CanoScan LiDE 200 scanner. Background obtained with blue paper sheets and a 30 cm ruler (Helix) fixed on the margins.

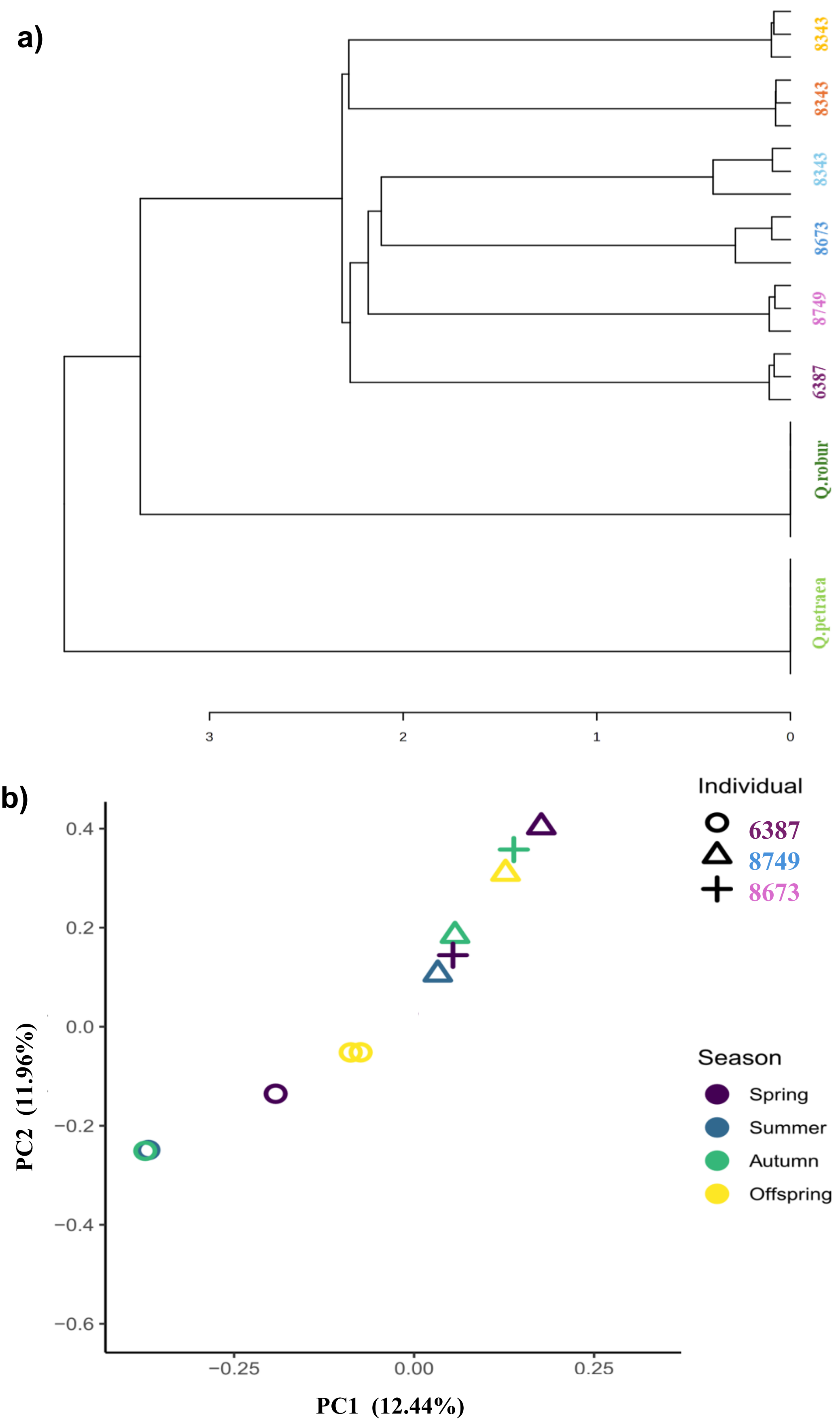


**Figure Supplementary 2. Genetic diversity among individuals. a)** Genotyping by sequencing of the 6 trees analysed with control samples of pure *Q.robur* and *Q.petraea*. Hierarchical clustering using Pearson, average method is represented. **b)** PCA plot of the genetic covariance between families and individuals (ST1) based on C/T and G/A filtered SNPs is represented. Shapes indicate the individual/ family of samples, and the colour indicated the season of collection.

a)

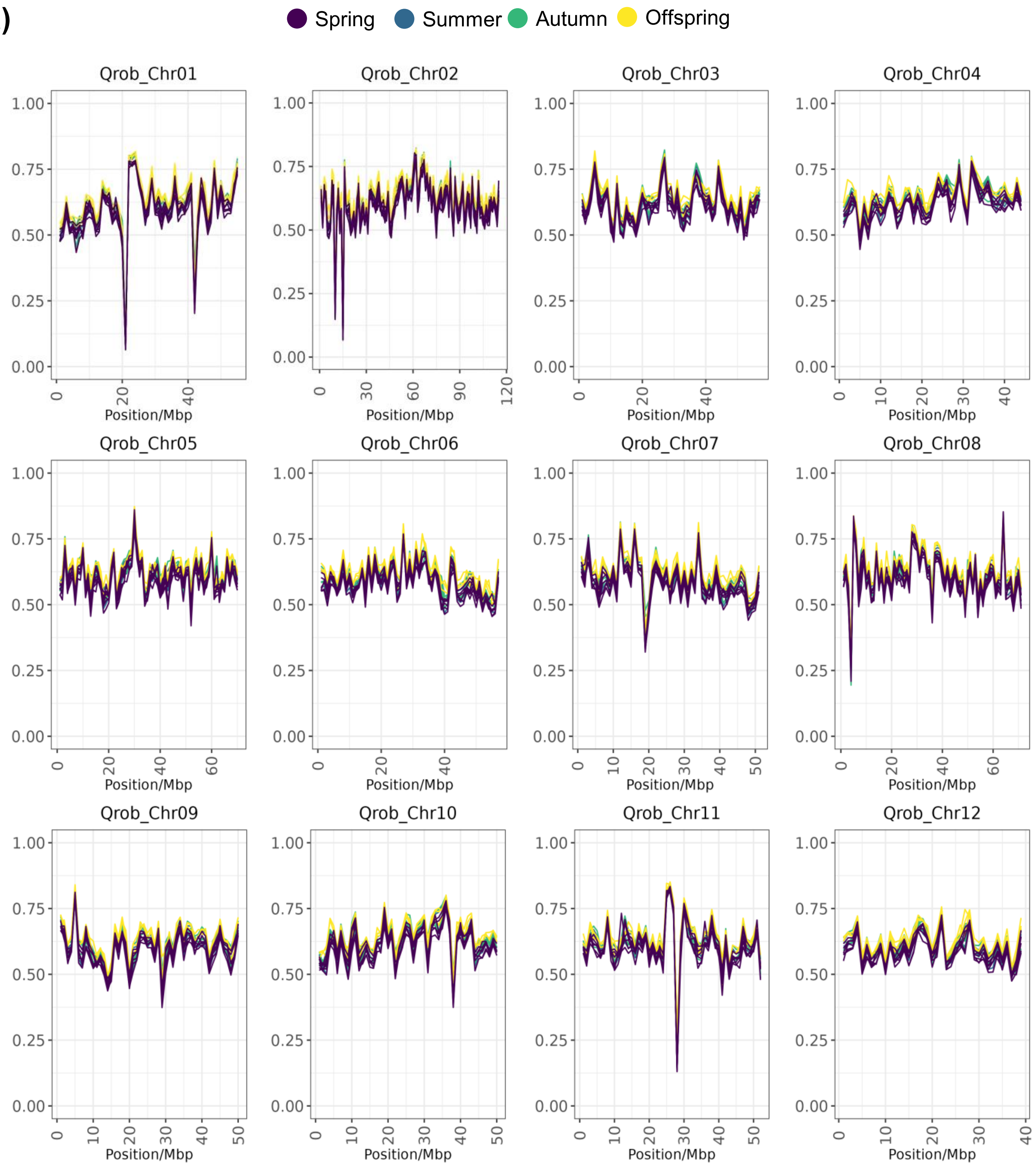


**Figure Supplementary 3: Methylation profiles by chromosome and context (a) CG, (b) CHG and (c) CHH.**

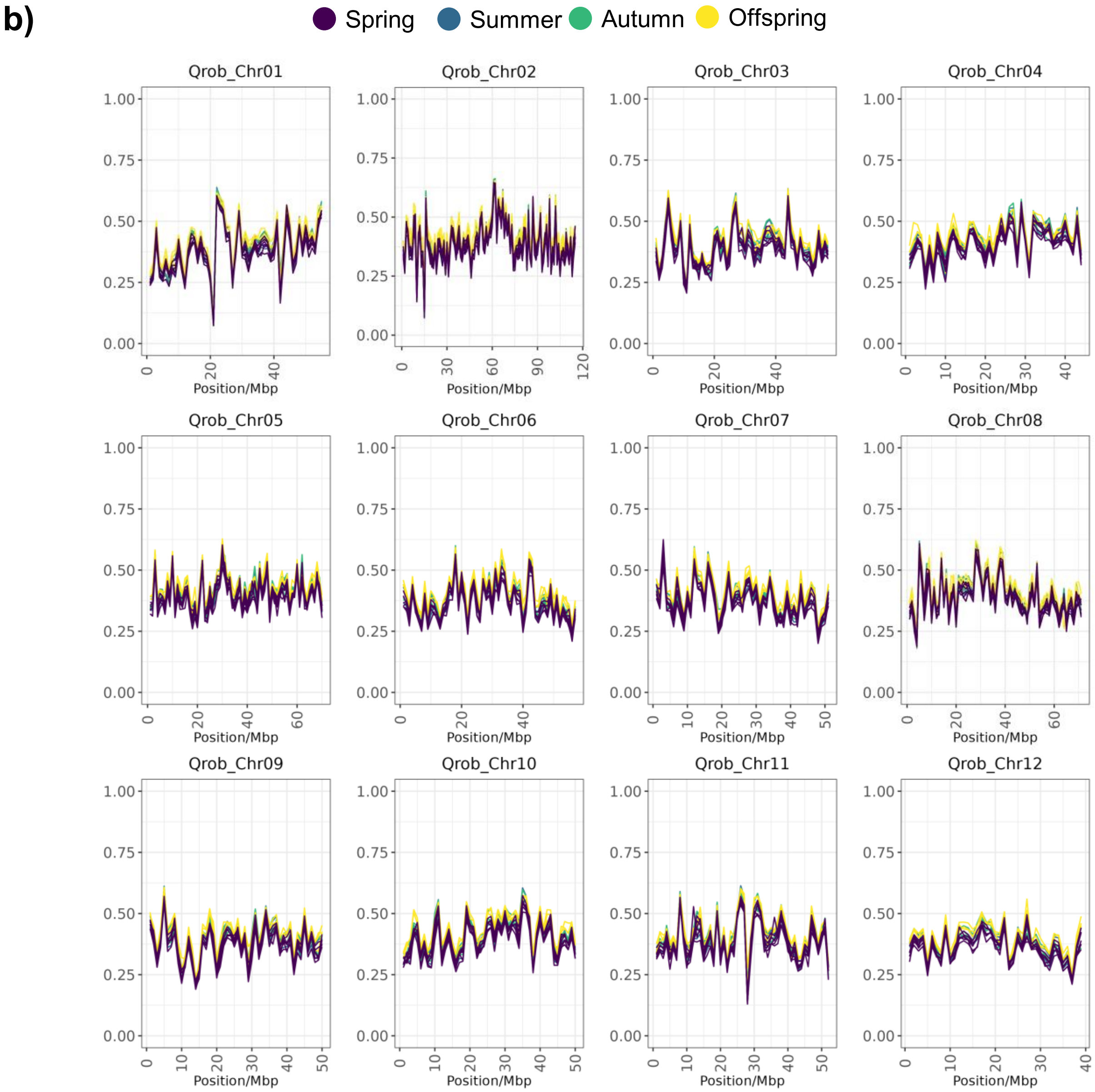


**Figure Supplementary 3 (continuation)**

c)

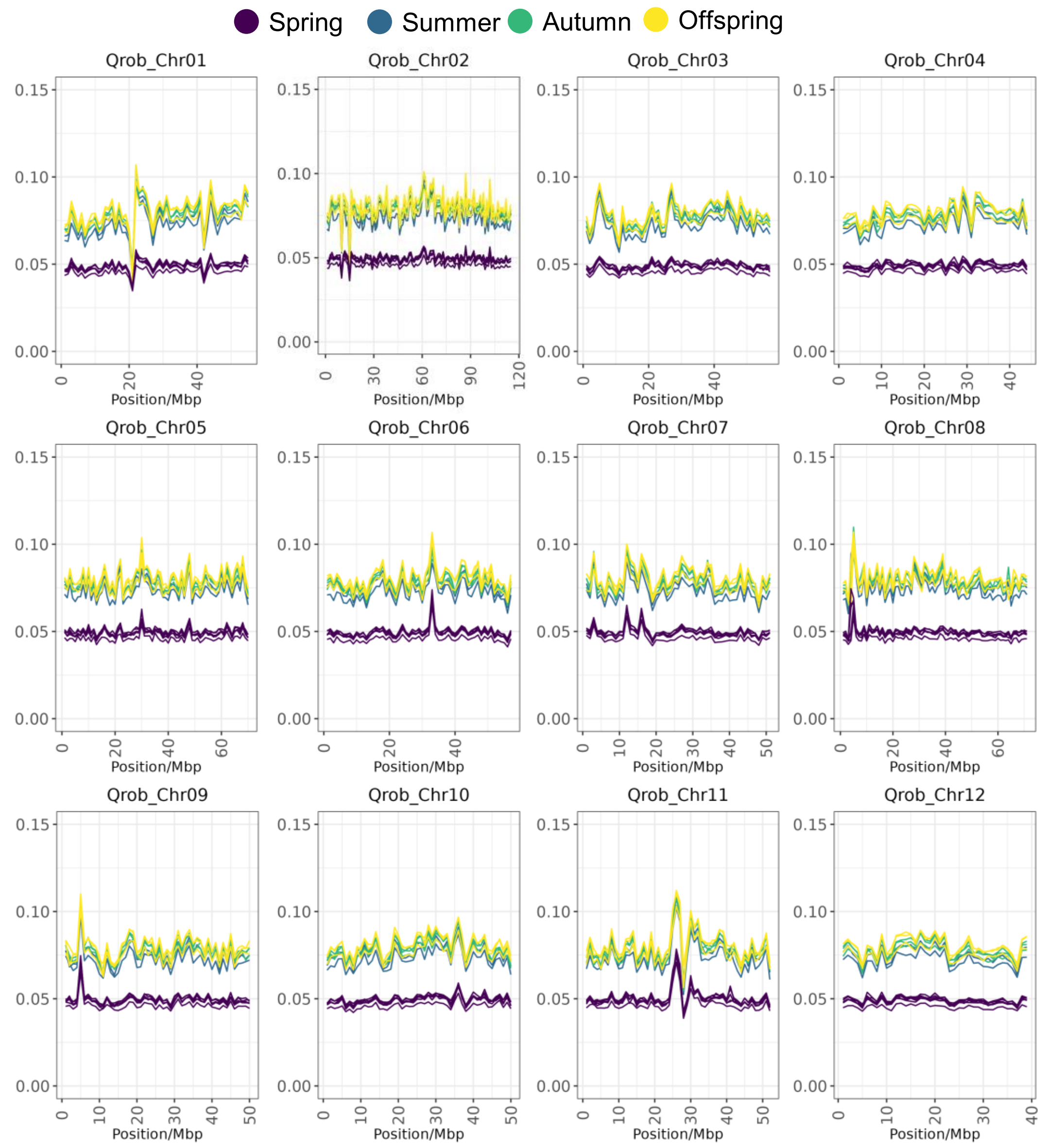


**Figure Supplementary 3 (continuation)**

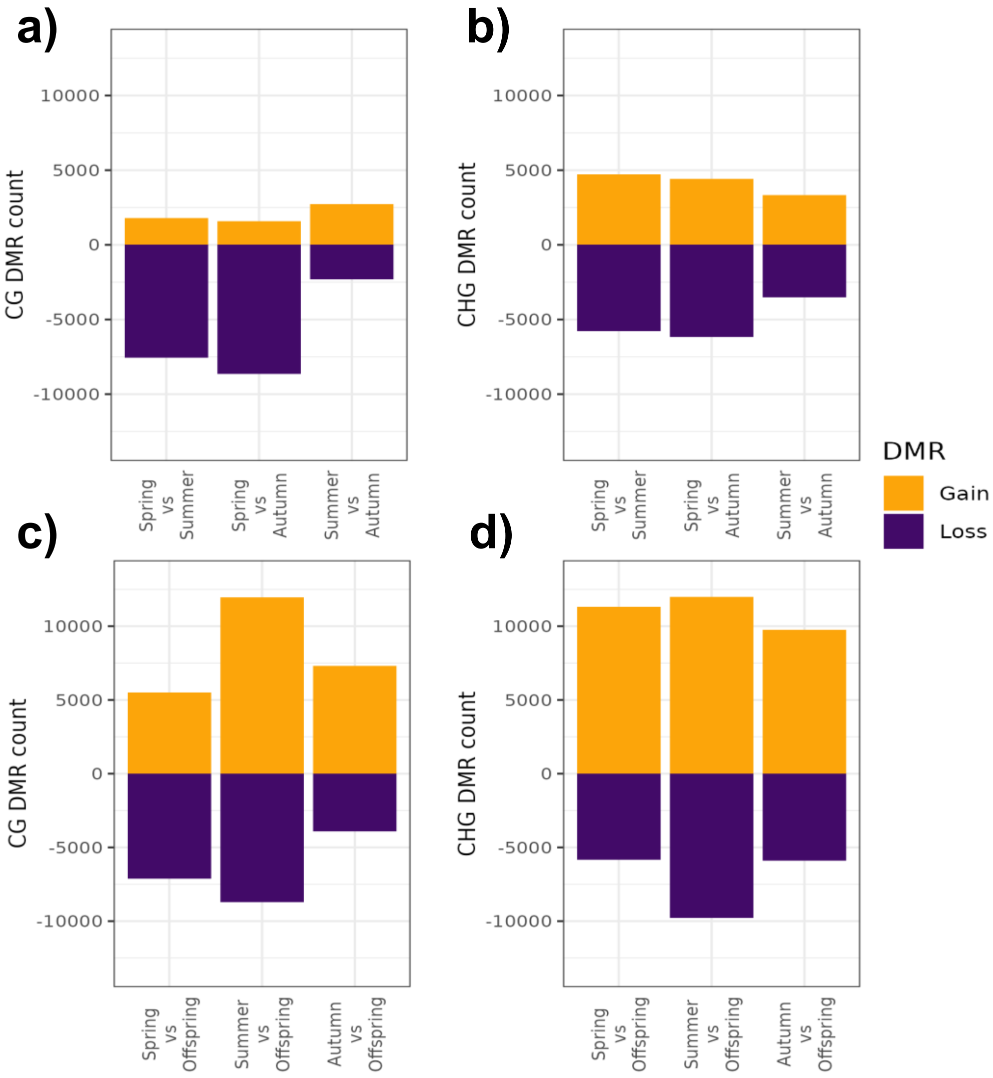


**Figure Supplementary 4. Number of total DMRs found between seasons and generations in the CG and CHG**. Seasonal DMR patterns in CG context (**a**) and CHG context (**b**) are represented together with the generational DMR patterns in CG (**c**) and CHG (**d**) contexts. Colours indicate hypermethylation (orange) or hypomethylation (blue) among the different comparisons.

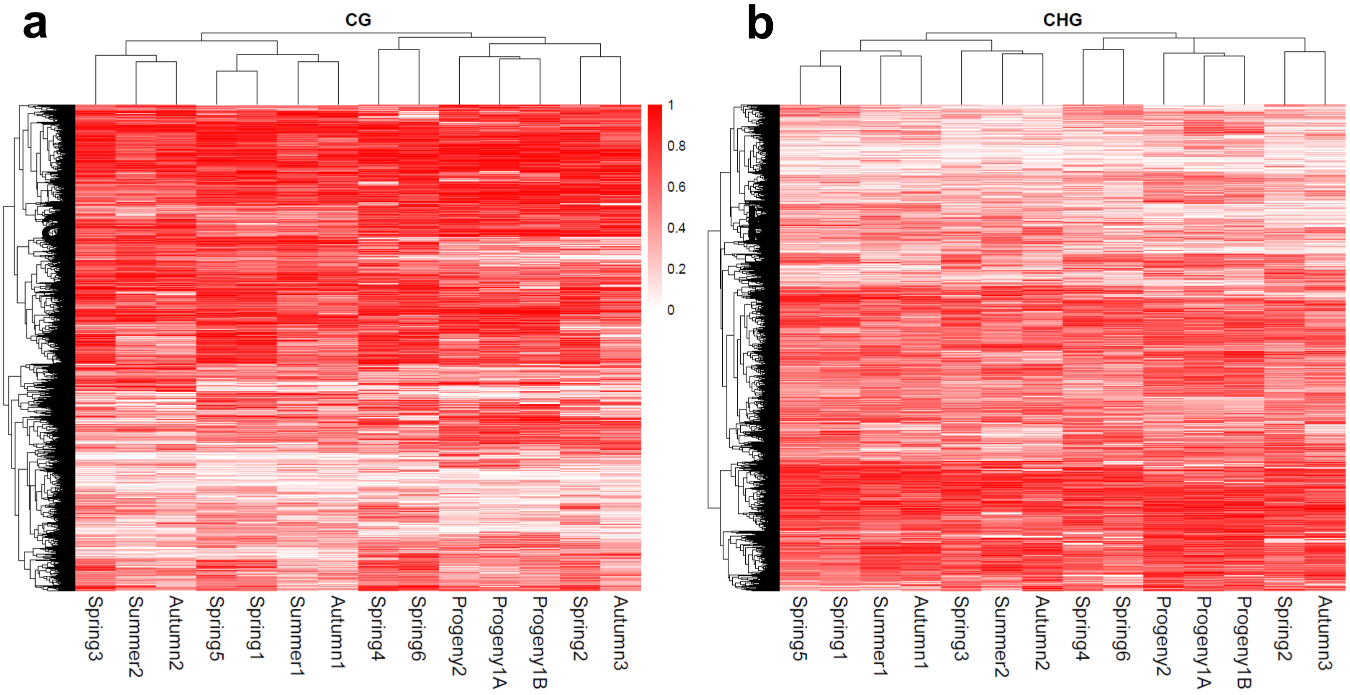


**Figure Supplementary 5. Methylation on DMRs loci,** Heatmaps of the methylation status **in** the CG (**a**) and CHG (**b**) contexts are represented.

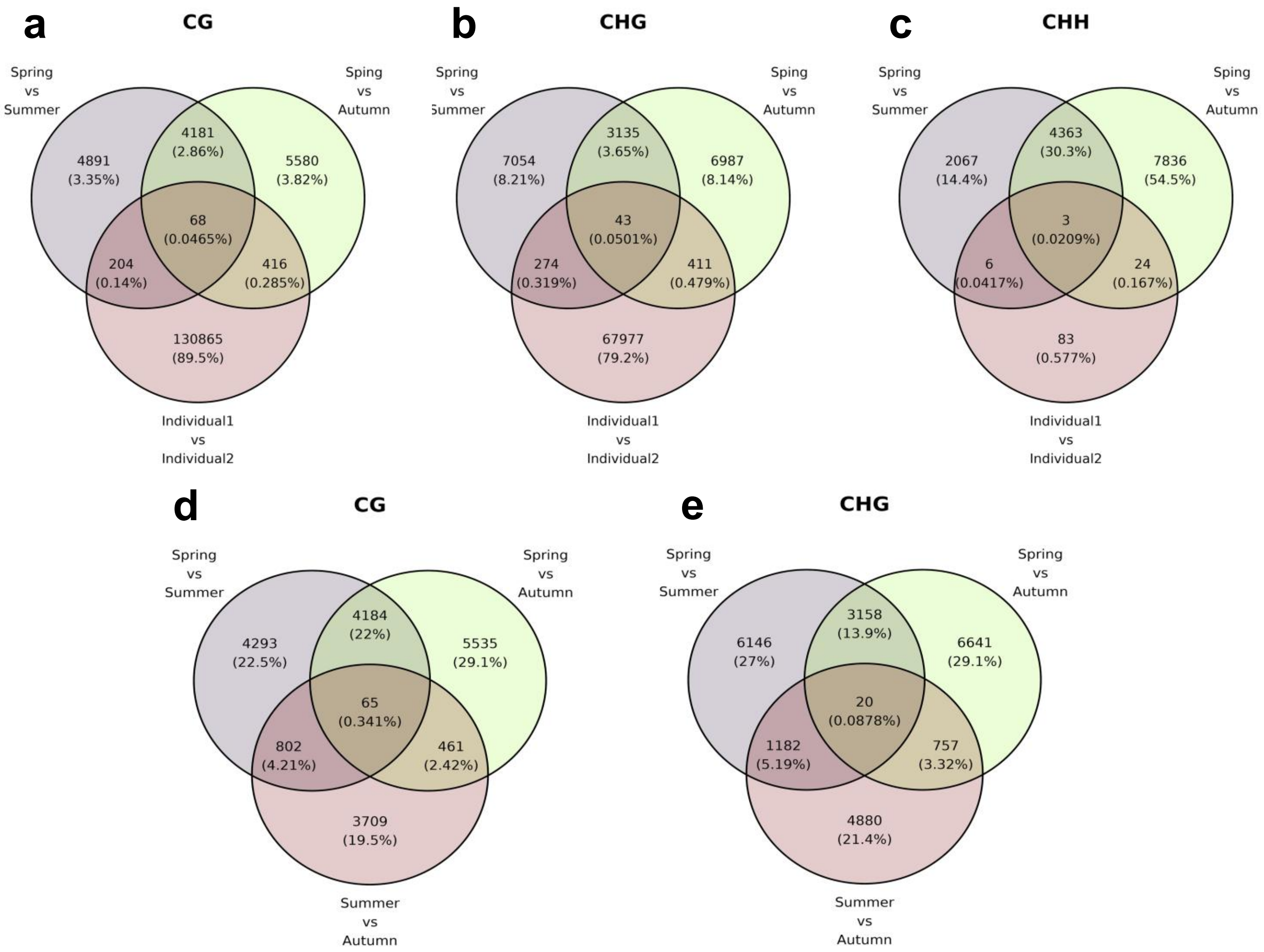


**Figure Supplementary 6. DMR overlaps between individuals and among seasons.** DMRs called from Individual 1 and Individual 2 samples only and the Spring Seasonal DMRs at the context CG **(a)**, CHG **(b)** and CHH **(c)**. DMRs in different seasons are presented in the CG **(d)** and CHG contexts **(e).**

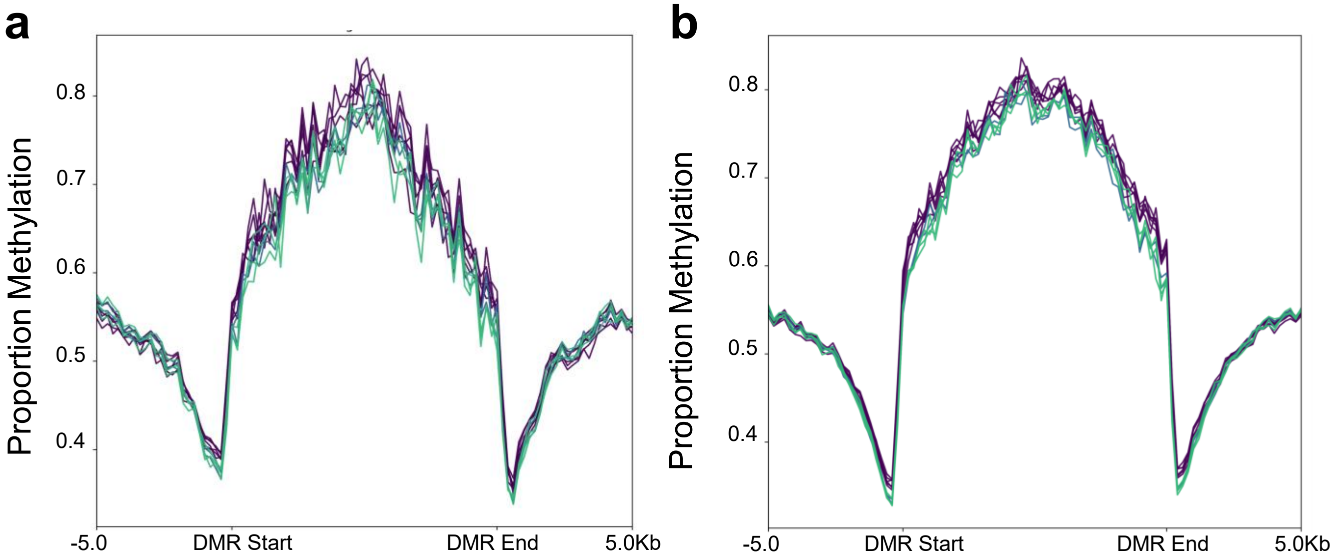


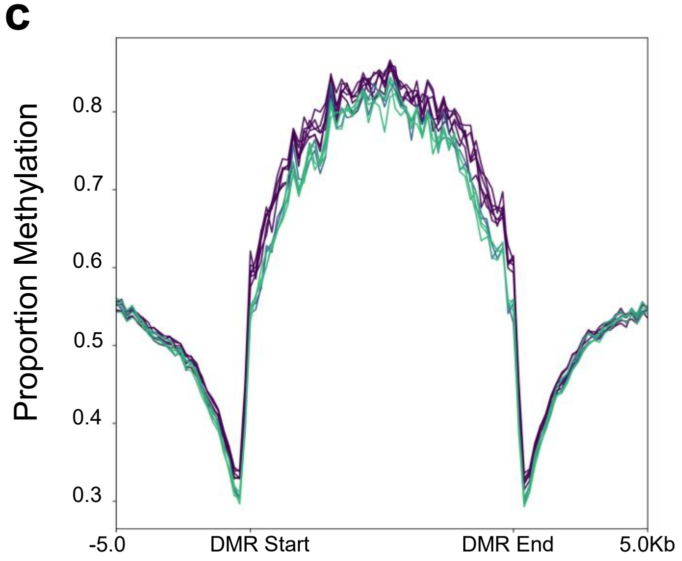


**Figure Supplementary 7. CG Methylation at CHH DMR loci by Intersection of Seasonal DMR calls**. CG methylation of Spring Seasonal CHH DMRs for the Spring/Summer only **(a),** Spring/Autumn only **(b)** and the intersect of Spring/Summer and Spring Autumn **(c)** are represented.

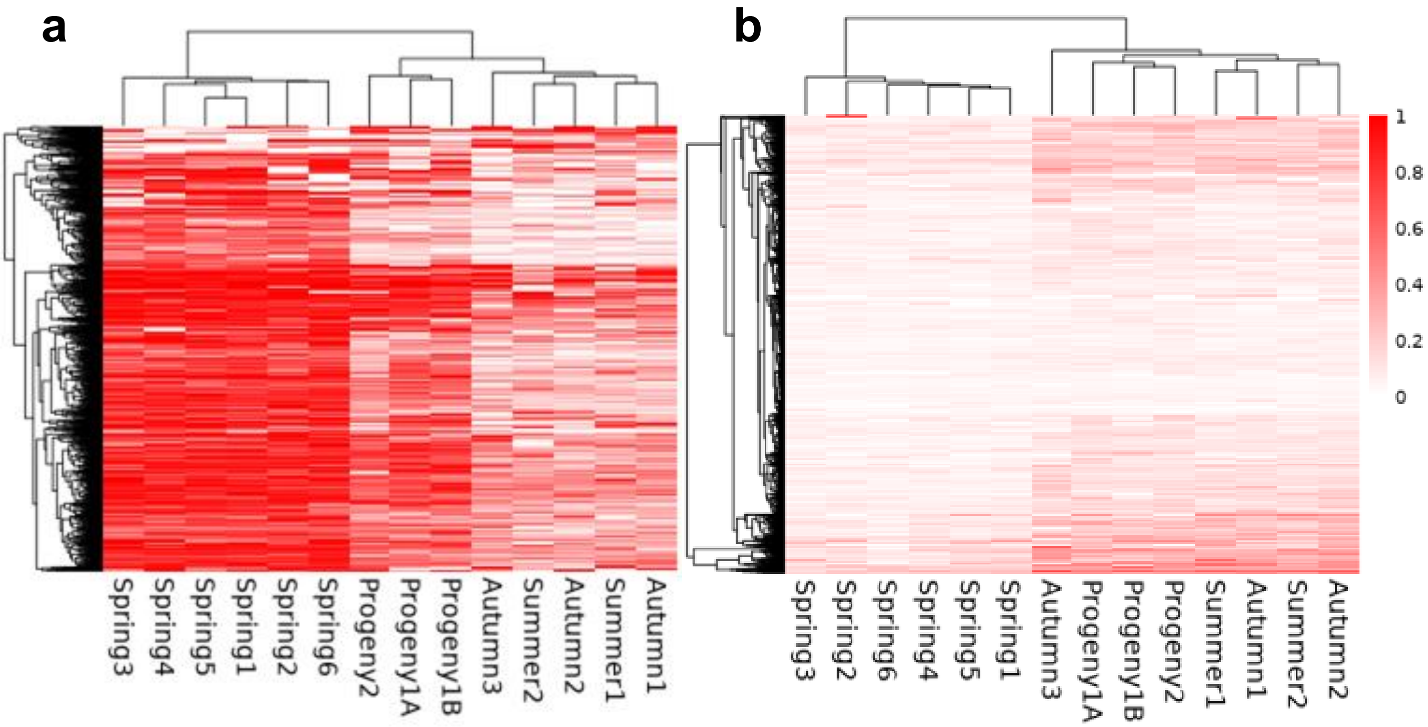


**Figure Supplementary 8. CG Methylation in Highly Differentially Methylated CHH DMR Loci.** The most highly differentially methylated (> 0.4) DMRs in CHH were represented **(a)** (n=2855) together with the same regions on **(b)** CG context (n = 386).

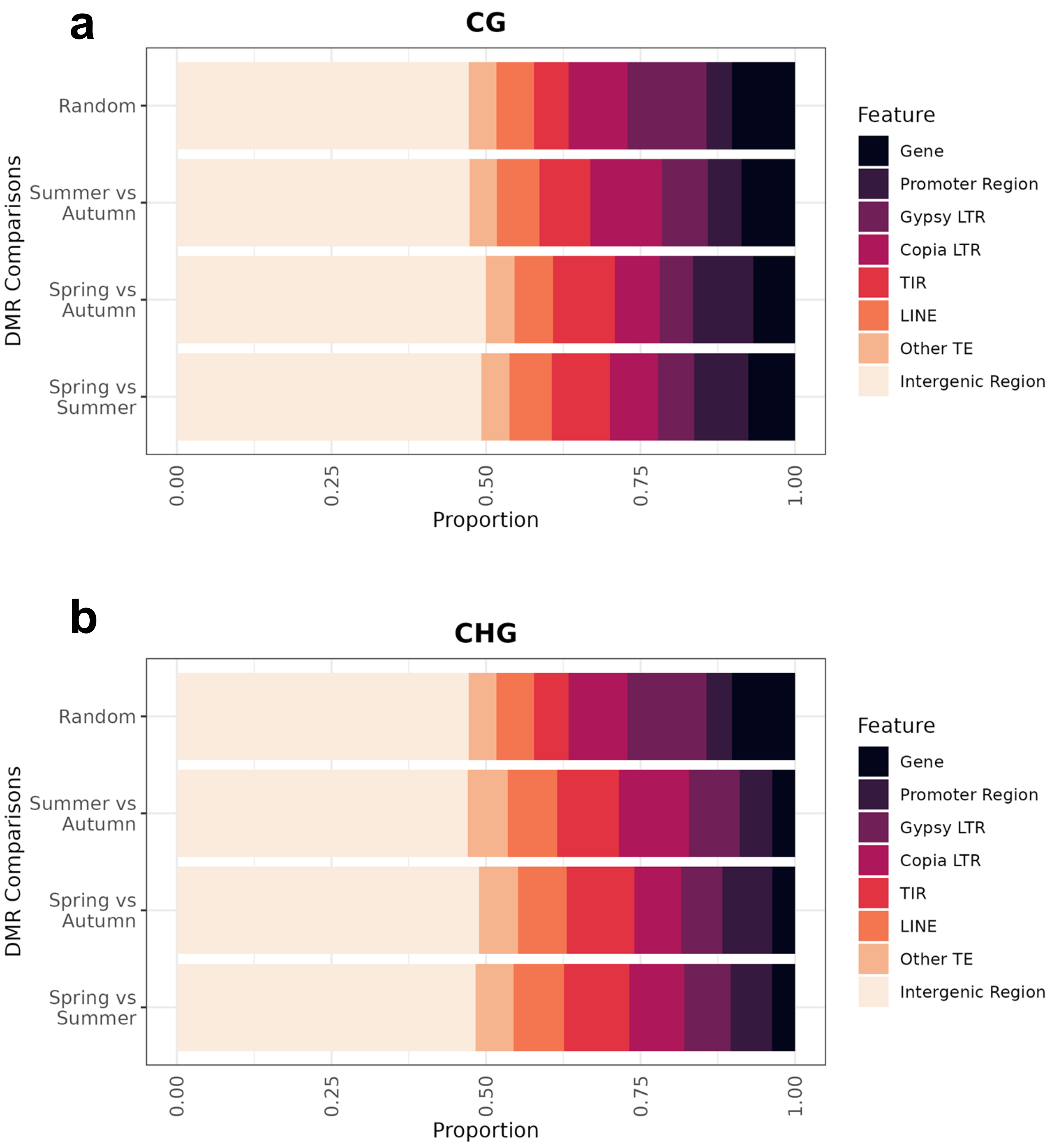


**Figure Supplementary 9. DMR Overlaps Between Seasons in CG and CHG Contexts** . DMRs were called between seasons and overlapped with genetic elements from the reference oak genome in the CG **(a)** and CHG **(b)** contexts. A “Random” bar under a null hypothesis where DMRs are placed randomly is simulated for comparison.

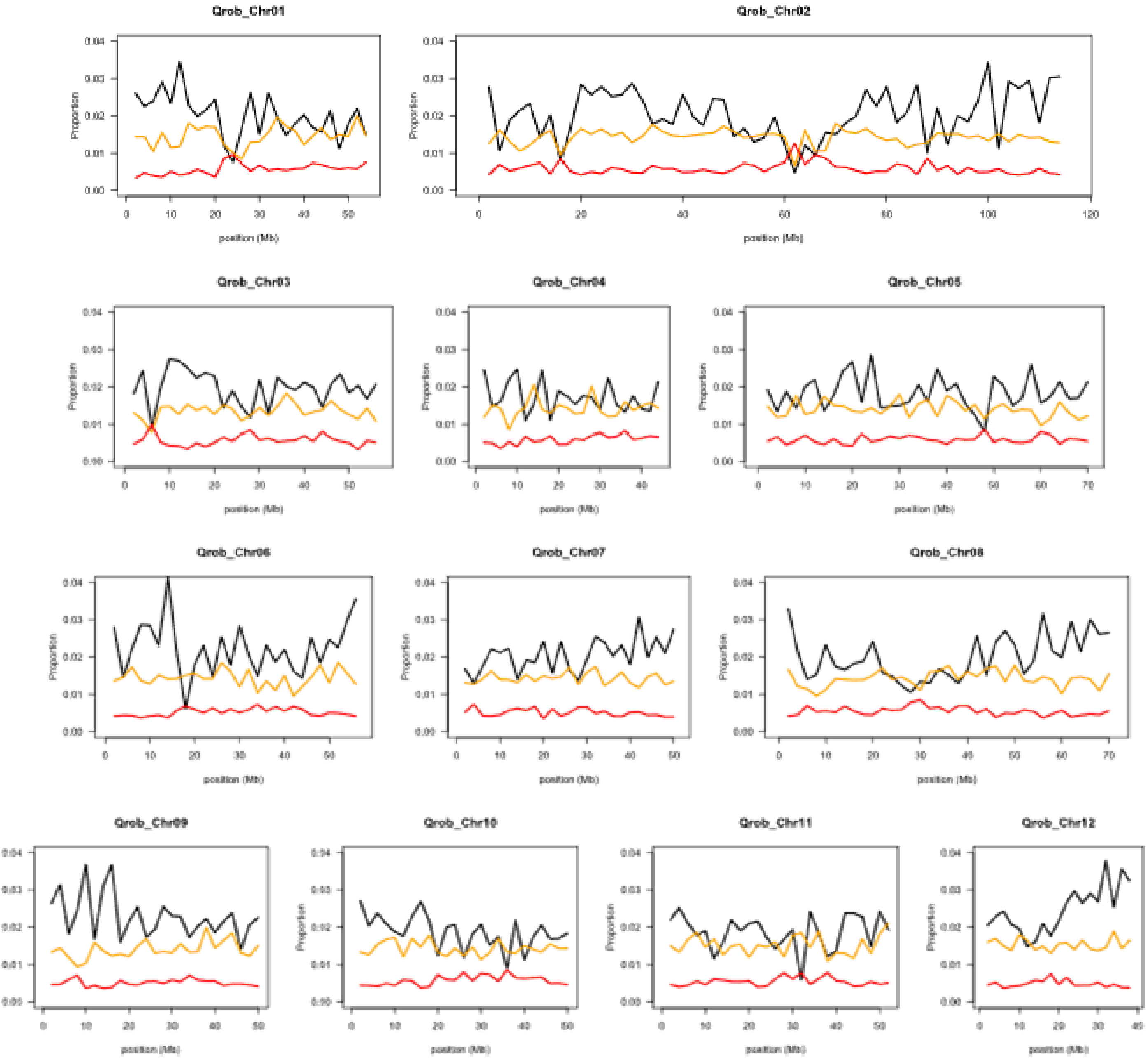


**Figure Supplementary 10**. **Gene and TE features distribution in Oak genome.** For annotated Genes (**black line**), TIR-TEs (**orange lines**) and LTR-TE (**red lines**), the densities have been calculated for each 200 Kb genomic tile and plotted for the 12 oak chromosomes.

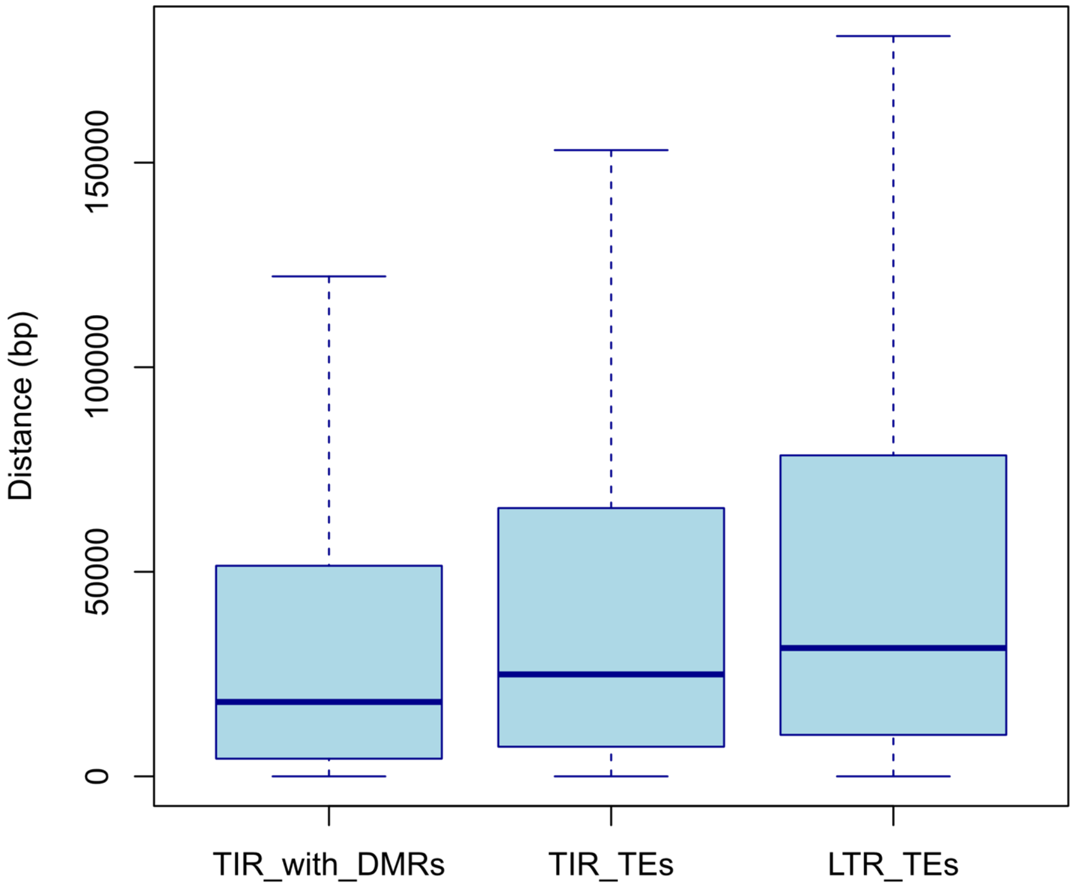


**Figure Supplementary 11. Distance from TEs to Nearest Genes. For every TIRs overlapping with DMRs.**

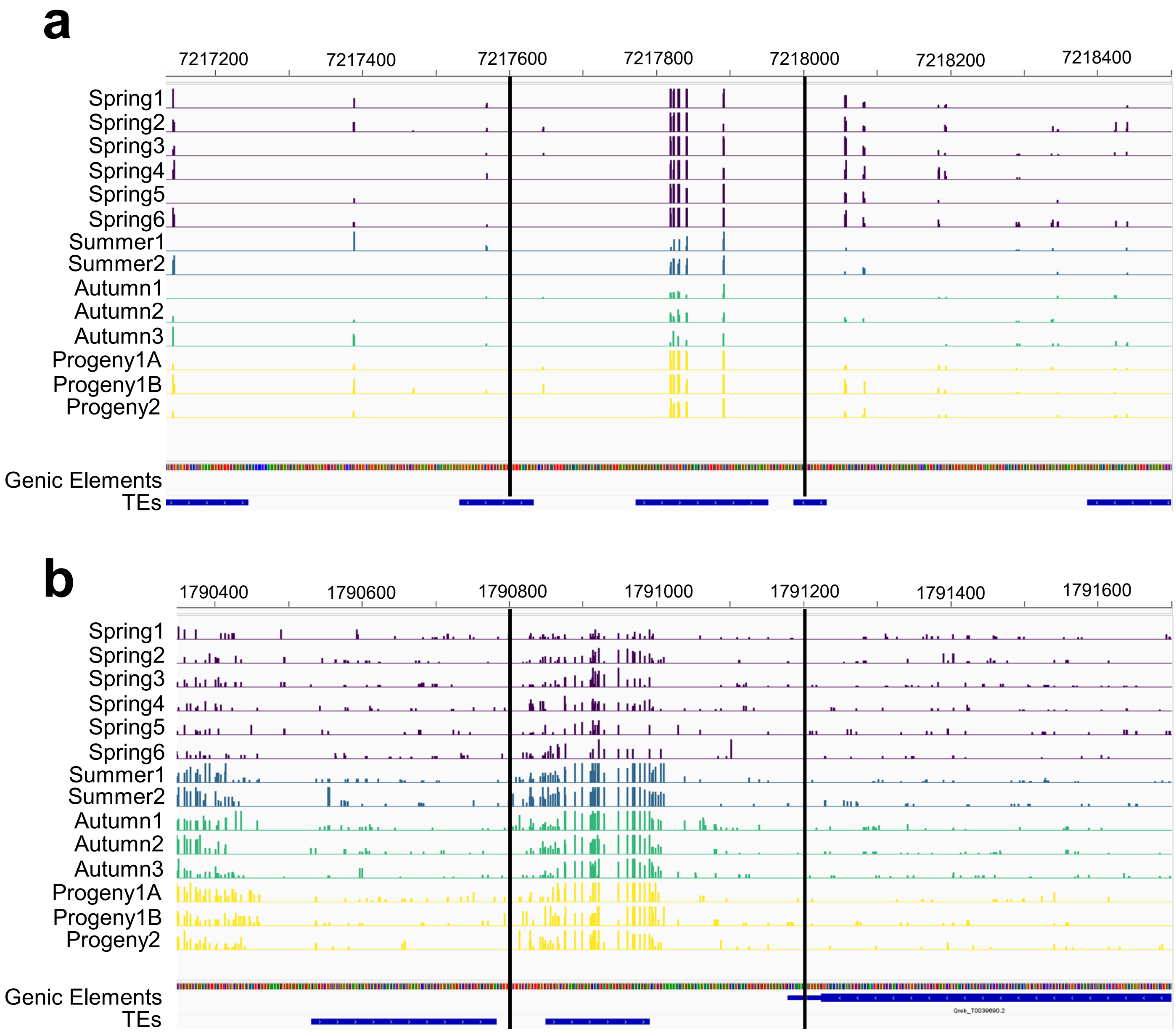


**Figure Supplementary 12. Examples of DMRs overlaps among seasons and off-springs**. **(a)** Example of a DMR identified (between vertical lines) in the CG context that overlaps between each of the 3 Seasonal comparisons and between Parents and Progeny (Qrob_Chr01 position 7217601-7218000). **(b)** Example of a DMR identified (between vertical lines) in the CHH context between Spring and Summer (Qrob_Chr01 position 1790801-1791200). Alignments are made with respect to the reference genome and annotations are associated with the same genome. Each track indicates the proportion of methylation scaled between 0-1.
